# Functional characterization of clinical isolates of the opportunistic fungal pathogen *Aspergillus nidulans*

**DOI:** 10.1101/2020.01.28.917278

**Authors:** Rafael Wesley Bastos, Clara Valero, Lilian Pereira Silva, Taylor Schoen, Milton Drott, Verônica Brauer, Rafael Silva-Rocha, Abigail Lind, Jacob L. Steenwyk, Antonis Rokas, Fernando Rodrigues, Agustin Resendiz-Sharpe, Katrien Lagrou, Marina Marcet-Houben, Toni Gabaldón, Erin McDonnell, Ian Reid, Adrian Tsang, Berl R. Oakley, Flávia Vieira Loures, Fausto Almeida, Anna Huttenlocher, Nancy P. Keller, Laure Nicolas Annick Ries, Gustavo H. Goldmana

## Abstract

*Aspergillus nidulans* is an opportunistic fungal pathogen in patients with immunodeficiency and virulence of *A. nidulans* isolates has mainly been studied in the context of the chronic granulomatous disease (CGD), with characterization of clinical isolates obtained from non-CGD patients remaining elusive. This study therefore carried out a detailed biological characterization of two *A. nidulans* clinical isolates (CIs), obtained from a patient with breast carcinoma and pneumonia and from a patient with cystic fibrosis that underwent lung transplantation, and compared them to the reference, non-clinical A4 strain. Both CIs presented increased growth in comparison to the reference strain in the presence of physiologically-relevant carbon sources. Metabolomic analyses showed that the three strains are metabolically very different from each other in these carbon sources. Furthermore, the CIs were highly susceptible to cell wall perturbing agents but not to other physiologically-relevant stresses. Genome analyses identified several frame-shift variants in genes encoding cell wall integrity (CWI) signalling components. Significant differences in CWI signalling were confirmed by western blotting among the three strains. *In vivo* virulence studies using several different models revealed that strain MO80069 had significantly higher virulence in hosts with impaired neutrophil function when compared to the other strains. In summary, this study presents detailed biological characterization of two *A. nidulans sensu stricto* clinical isolates. Just like in *A. fumigatus,* strain heterogeneity exists in *A. nidulans* clinical strains that can define virulence traits. Further studies are required to fully characterize *A. nidulans* strain-specific virulence traits and pathogenicity.

**Importance:** Immunocompromised patients are susceptible to infections with opportunistic filamentous fungi from the genus *Aspergillus.* Although *A. fumigatus* is the main etiological agent of *Aspergillus* spp.-related infections, other species, such as *A. nidulans* are prevalent in a condition-specific manner. *A. nidulans* is a predominant infective agent in patients suffering from chronic granulomatous disease (CGD). *A. nidulans* isolates have mainly been studied in the context of CGD, although infection with *A. nidulans* also occurs in non-CGD patients. This study carried out a detailed biological characterization of two non-CGD *A. nidulans* clinical isolates and compared it to a reference strain. Phenotypic, metabolomic and genomic analyses highlight fundamental differences in carbon source utilization, stress responses and maintenance of cell wall integrity among the strains. One clinical strain had increased virulence in models with impaired neutrophil function. Just as in *A. fumigatus,* strain heterogeneity exists in *A. nidulans* clinical strains that can define virulence traits.

## Introduction

Fungal pathogen-related infections are now estimated to result in a higher number of human deaths as tuberculosis or malaria alone (1–3). The majority of systemic fungal infections are caused by *Candida spp., Pneumocystis* spp., *Cryptococcus spp.* and *Aspergillus spp.* (4, 5). Of the hundreds of known *Aspergillus spp.,* only a few cause disease in animals, with the most prominent being *A. fumigatus, A. flavus, A. nidulans, A. niger* and *A. terreus* (6, 7).

The primary route of infection of *Aspergillus spp.* is via the inhalation of conidia (asexual spores). In immunocompetent individuals, inhaled conidia are rapidly cleared by pulmonary resident and recruited neutrophils and macrophages, together preventing the onset of infection (8–10). However, disturbances to the immune system may render an individual susceptible to infection by *Aspergillus spp.* (11). The severity of infection largely depends on fungal species and genotype, the host immunological status and host lung structure (6). Invasive Aspergillosis (IA) is the most severe disease caused by *Aspergillus spp.,* and is characterized by systemic host invasion, resulting in high mortality rates (30-95%) (2,10,11).

Patient populations with the highest risk of IA are those with prolonged neutropenia from intensive myeloablative chemotherapy and those with genetic disorders resulting in primary immune deficiencies, such as chronic granulomatous disease (CGD) (12, 13). CGD is a genetic disorder that affects 1 in 250,000 people and in ~80% of all cases subjects are of the male sex. CGD is caused by mutations in the genes encoding any of the five structural components of the Nicotinamide Adenine Dinucleotide Phosphate (NADPH) - oxidase complex, an enzyme complex important for superoxide anion and downstream reactive oxygen species (ROS) production in phagocytic cells (14). As a result, immune cells are unable to efficiently kill microorganisms and these microorganisms can then become pathogenic in such patients (13, 14)

Although *A. fumigatus* is the main etiological agent of *Aspergillus-related* infections in immunocompromised patients; other *Aspergillus* spp. have been found to have a high infection rate under some conditions. *A. nidulans* infections are not commonly reported in immunocompromised patients, except for subjects suffering from CGD (15, 16). In CGD patients, *A. fumigatus* and *A. nidulans* are responsible for 44% and 23% respectively, of all fungal infections (15, 16). Infections with *A. nidulans* cause mortality in 27-32% of CGD patients (15) and in comparison to *A. fumigatus, A. nidulans* isolates have higher virulence, invasiveness and dissemination, and resistance to antifungal drugs in these patients (17). Hence, *A. nidulans* infections have been studied mainly in the context of CGD although this fungal species can also be virulent in non-CGD, immunocompromised patients (18). In comparison to *A. fumigatus,* investigations into *A. nidulans* isolate virulence have been neglected with very few studies having investigated the genetic and metabolic features of *A. nidulans* clinical strains, isolated from CGD and non-CGD patients, in the context of stress responses encountered during human host infection as well as when interacting with host immune responses (18–21).

The aim of this work was to carry out a detailed molecular, phenotypic and virulence characterization of two *A. nidulans* clinical isolates from a) a patient with breast carcinoma and pneumonia and b) a patient with cystic fibrosis who underwent lung transplantation and compare them to the well-characterized, wild-type isolate FGSC A4.

## Results

### The *A. nidulans* clinical isolates have increased growth, in comparison to the reference strain, in the presence of alternative carbon sources

Fungal metabolic plasticity, which allows growth in unique and diverse ambient and host microenvironments, has long been hypothesized to contribute to *Aspergillus* virulence, with carbon sources such as glucose (22), ethanol (23) and acetate (24) being predicted to be actively used during *in vivo* infection. In addition, fatty acids and lipids are also thought to serve as major nutrient sources during mammalian host colonization as is evident by the importance of key glyoxylate cycle enzymes in fungal virulence (25). We therefore characterized growth, by determining fungal dry weight, of the two *A. nidulans* CIs in the presence of minimal medium (MM) supplemented with different physiologically-relevant carbon sources, namely glucose, acetate, ethanol and lipids, and compared it to the FGSC A4 reference strain. A significant reduction in growth was observed for both CIs in the presence of glucose whereas they had significantly increased growth in the presence of the alternative carbon source ethanol, casamino acid and the lipids Tween 20 (a source of lauric, palmitic, and myristic acids) (26), Tween 80 (which contains principally oleate) (26) and olive oil (triacylglycerols and free fatty acids) (27) (Fig. 1). In contrast, no difference in fungal biomass accumulation was observed in the presence of acetate and the lung-resident glycoprotein mucin (Fig. 1). These results suggest that the *A. nidulans* CIs have improved growth relative to the reference strain in the presence of most of the alternative carbon sources tested here, including different lipids.

**Figure 1.**
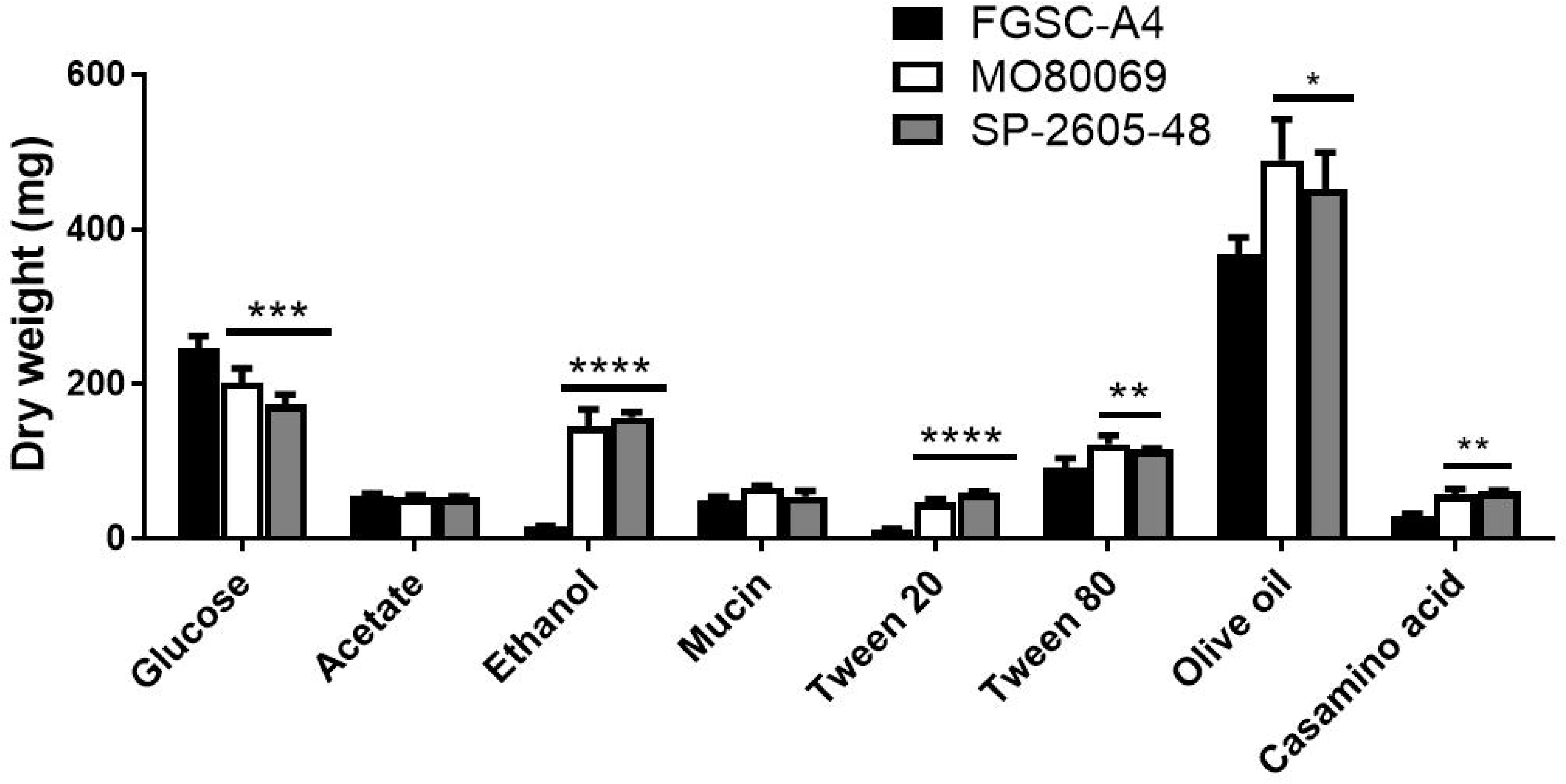
The *A. nidulans* clinical isolates exhibit improved growth in the presence of alternative carbon and lipid sources. Strains were grown in liquid MM supplemented with glucose, acetate, ethanol, mucin, tween 20 and 80, olive oil and casamino acids at 37°C for 48 h (glucose) or 72 h (others) before fungal biomass was freeze-dried and weighed. Standard deviations were determined from biological triplicates with \*\**p*<0.01; \*\*\**p*<0.001; \*\*\*\**p*<0.0001 in a one-way ANOVA with Tukey post-test comparing growth of the clinical isolates to the FGSC-A4 reference strain.

### Metabolic profiles differ among the *A. nidulans* clinical isolates and reference strain in the presence of different carbon sources

To further investigate nutrient utilization in the *A. nidulans* CIs, the metabolic profiles of strains MO80069 and SP-2605-48 were determined and compared to the reference strain A4. Metabolomics was carried out on cellular extracts from strains grown for 24 h in fructose-rich MM and then transferred for 16 h to MM supplemented with glucose (CIs present reduced growth), ethanol (CIs had increased growth), acetate and mucin (no difference in growth). A total of 40 different metabolites were identified when strains were grown in the presence of glucose and ethanol, whereas 44 different metabolites were identified when strains were grown in the presence of acetate and mucin (Table S1). When comparing metabolite quantities of strain MO80069 to the reference strain, 18 (45%), 22 (55%), 23 (52%) and 24 (55%) metabolite quantities were significantly (p-value < 0.05) different from the quantities in the reference strain when grown in glucose, ethanol, acetate and mucin respectively (Table S1, Table 1). In strain SP-2505-48, 15 (38%), 23 (58%), 30 (68%) and 14 (32%) metabolite quantities, that were normalized by fungal dry weight, were significantly (p-value < 0.05) different from the quantities in the reference strain in the presence of glucose, ethanol, acetate and mucin respectively (Table S1, Table 1). Principal component analysis (PCA) and Hierarchical clustering analysis (HCA) of identified metabolite quantities showed that the CIs clustered apart from the reference strain and from each other in all tested carbon sources (Fig. S1-S2), indicating that they are metabolically different from the reference strain and from each other.

**Table 1.**
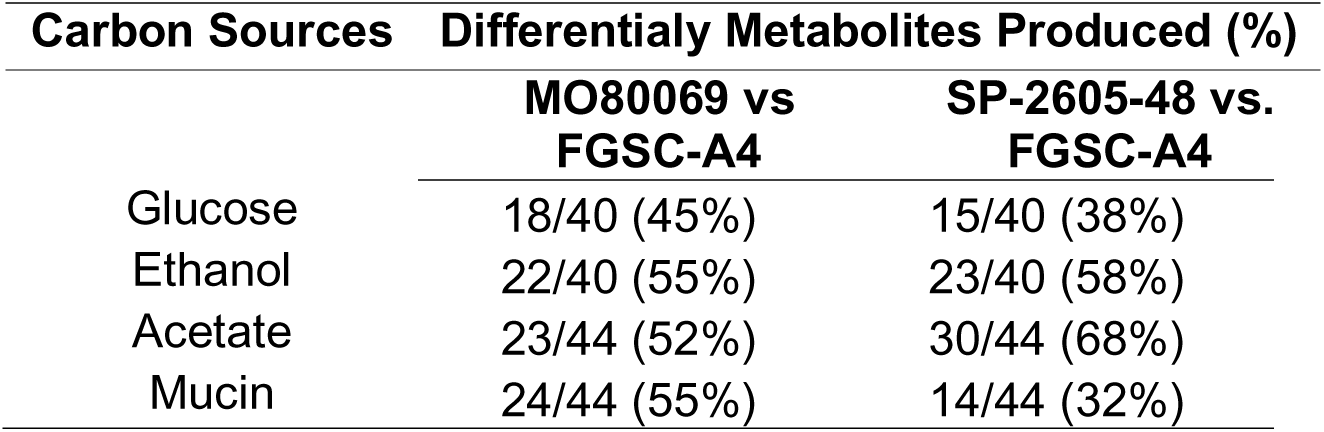
Number and percentage of identified metabolite quantities that were significantly (*p*-value < 0.05) different in the *A. nidulans* clinical isolates in comparison to the reference strain when strains were grown in the presence of glucose, ethanol, acetate and mucin for 16 h.

When further focusing on metabolites that were significantly different in quantity between the CIs and the reference strain, we observed that in the presence of glucose and ethanol, the majority of identified metabolites were present in significant lower quantities in comparison with the reference strain; whereas both CIs had significant higher metabolite quantities in the presence of acetate in comparison with the reference strain (Fig. 2A-C). Furthermore, when the *A. nidulans* CIs were cultivated in mucin-rich minimal medium, only 9 out of 29 significantly different metabolite quantities were identified in both strains whereas the remaining metabolite quantities were strain-specific, suggesting that the metabolic profiles of the two differed drastically the presence of this carbon source (Fig. 2D).

**Figure 2.**
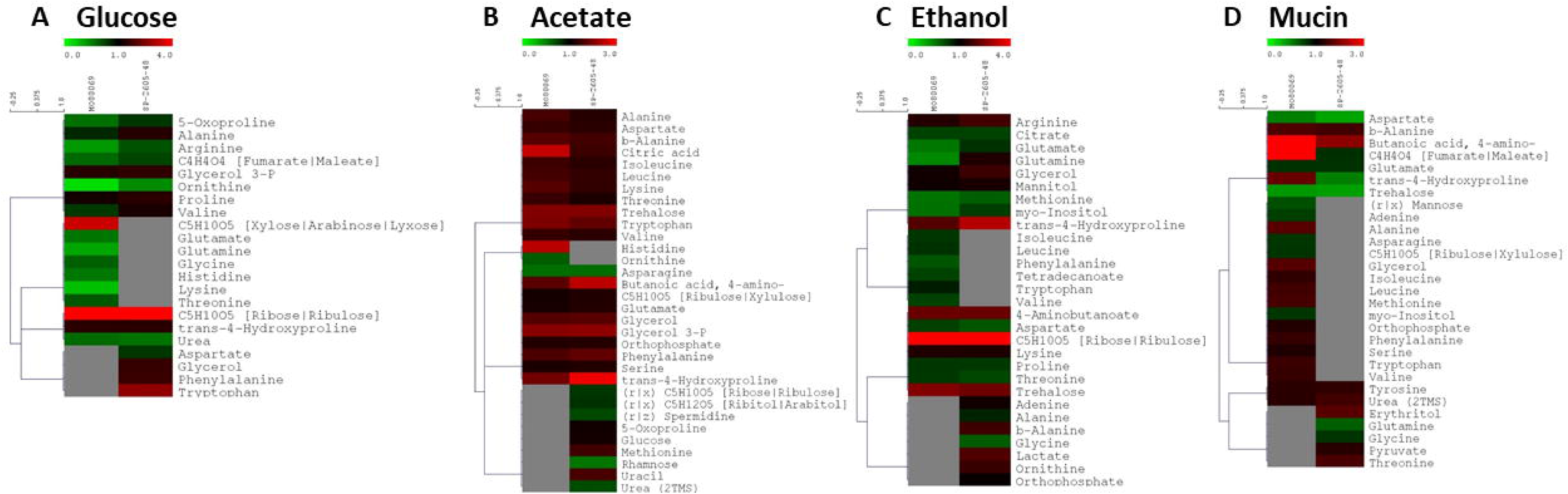
The *A. nidulans* clinical isolates are metabolically different from the reference strain in the presence of different carbon sources. Heat maps depicting log-fold changes of identified metabolite quantities, that were significantly (*p*<0.05) different in the *A. nidulans* clinical isolates MO80069 and SP-2605-48 when compared to the FGSC-A4 reference strain (grey squares depict metabolite quantities that were not detected as significantly different in one of the clinical isolates).

When the CIs were grown in a glucose-rich MM, amino acids were found in lower quantities in both CIs when compared to the reference strain. In contrast, pentose phosphate pathway (PPP) intermediates, glycerol, glycerol derivatives and aromatic amino acids were detected in significantly higher quantities in this carbon source (Fig. 2A). In an ethanol-rich MM, significantly lower quantities of various amino acids as well as of the citric acid cycle intermediate citrate were detected in the CIs; whereas increased quantities of several amino acid pathway intermediates, the carbon compounds glycerol, mannitol and trehalose, PPP intermediates and lactate were detected in the CIs when compared to the reference strain in this carbon source (Fig. 2C). In acetate-rich MM, most identified metabolites, notably a variety of amino acids, were present in significantly higher amounts in the CIs when compared to the reference strain, with the exception of some amino acids, PPP intermediates, spermidine, rhamnose and urea (Fig. 2B). When strains were grown in mucin-rich MM, differences in the quantities of a variety of amino acids were observed, whereas trehalose was present in significantly lower quantities and urea in significantly higher quantities in both CIs when compared to the reference strain (Fig. 2D). In summary, these results suggest significant differences in amino acid biosynthesis and degradation, carbon source storage compounds and degradation among the different *A. nidulans* strains in a condition-dependent manner.

To determine if any metabolic pathways were specifically enriched in the *A. nidulans* CIs in comparison to the reference strain, pathway enrichment analyses was carried out on the metabolome data from glucose-, ethanol-, acetate- and mucin-grown cultures. In all tested carbon sources, with the exception of mucin for isolate SP-2605-48, there was significant enrichment for aminoacyl-tRNA biosynthesis (Table 2). The pathway constituting the metabolism of arginine and proline, was significantly enriched in both clinical isolates when grown in the presence of glucose and ethanol and in isolate SP-2605-48 when incubated in mucin-rich media (Table 2). When acetate was used as the sole carbon and energy source, enrichment of the metabolism of these amino acids was not observed (Table 2). In addition, metabolites identified for strain SP-2605-48 in the presence of mucin and ethanol showed pathway enrichment in nitrogen metabolism (Table 2). In agreement with the aforementioned differences in amino acid quantities, these results suggest that the Cis exhibit differences in nitrogen metabolism in a carbon source-independent manner when compared to the reference strain.

**Table 2.**
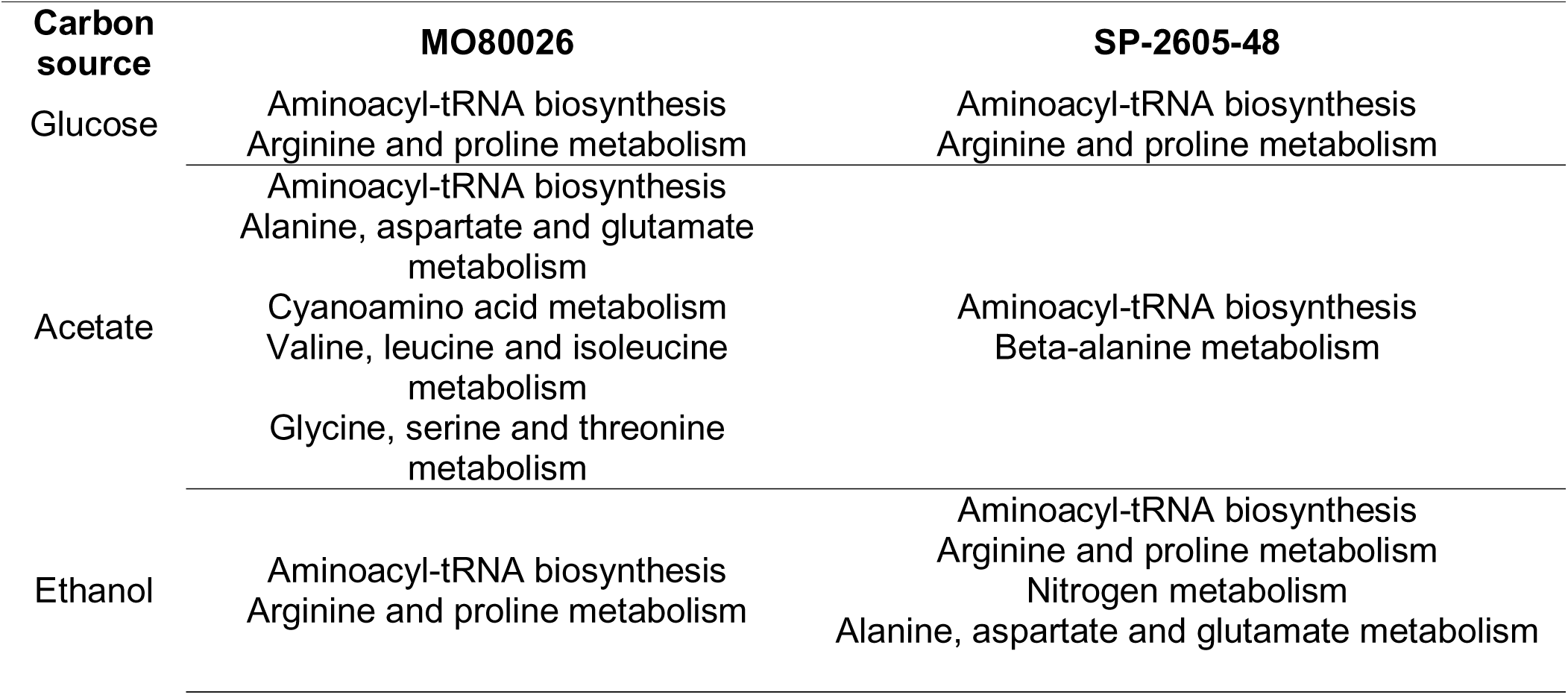
Significant metabolic pathway enrichments

### The *A. nidulans* clinical isolates are more sensitive to hydrogen peroxide-induced oxidative stress and cell wall-perturbing agents when compared to the reference strain

Due to the significant metabolic differences observed between the CIs and the reference strain in the presence of physiological-relevant carbon sources, and that primary metabolism (carbon source utilization) has been shown to impact virulence factors in opportunistic pathogenic fungi (28, 29), we hypothesized similar differences could be observed in the presence of physiological-relevant stress conditions. One such virulence factor is the fungal cell wall, which is crucial for protection, interaction with and modulation or evasion of the host immune system (30). In addition, cell wall polysaccharide composition is dependent on carbon source primary metabolism (28,29,31).

The production of reactive oxygen species (ROS), such as H_2_O_2_, and subsequent augmentation of cellular oxidative stress is a strategy employed by the mammalian immune system to combat potential invading pathogenic microorganisms (14). The *A. nidulans* reference strain and the two CIs were therefore grown in the presence of hydrogen peroxide (H_2_O_2_) and the oxidative stress-inducing compound menadione. Both CIs were more sensitive (reduced growth) to high concentrations of H_2_O_2_ (Fig. S3A), whereas they were resistant to menadione when compared to the reference strain (Fig. S3B). Furthermore, iron sequestration and elevated body temperature are additional physiological stress responses exerted by the host to prevent and/or control infection progression (32). Strains were therefore grown on iron-poor, glucose-rich minimal medium supplemented without (control) or with the iron chelators BPS and ferrozine (Fig. S3C), as well as in the presence of increasing temperatures (Fig. S3D). Growth of all strains was similar in these conditions, although strain MO80069 grew slightly more in the presence of the iron chelators (Fig. S3C). Lastly, growth of all strains was assessed in the presence of the cell wall perturbing agents caspofungin, congo red (CR) and calcofluor white (CFW). The echinocandin caspofungin is a competitive inhibitor of the cell wall enzyme β-1,3-glucan synthase (33) while CR and CFW bind to glucan or chitin chains respectively (34, 35). CR and CFW therefore interfere with the cross-linking of cell wall polysaccharides, resulting in a reduction of cell wall stability. Both clinical isolates were more sensitive to low and medium concentrations of caspofungin when compared to the reference strain, whereas all three strains grew similarly in the highest tested caspofungin concentration (8 μg/ml) (Fig. 3A). Similarly, both clinical strains were more sensitive to lower concentrations of CR whereas no significant difference in growth was observed in the presence of 50 μg/ml CR between all strains (Fig. 3B). In contrast, the CIs had significantly reduced growth in the presence of all tested CFW concentrations when compared to the reference strain (Fig. 3C).

**Figure 3.**
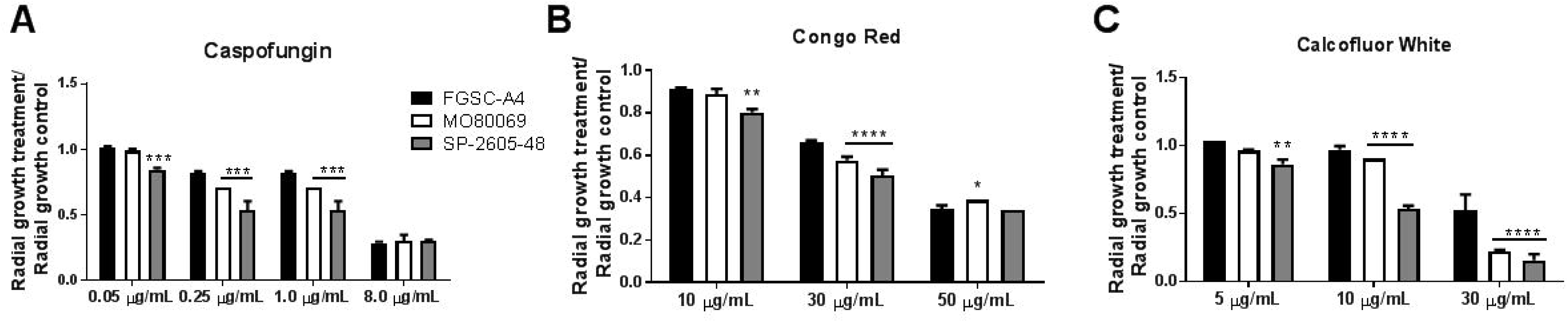
The A. nidulans clinical isolates are more sensitive to the cell wall-perturbing agents. Strains were grown from 105 spores on glucose minimal medium supplemented with increasing concentration of (A) caspofungin, (B) congo red and (C) calcofluor white for 5 days at 37°C. Standard deviations represent biological triplicates with \*\**p*<0.01; \*\*\**p*<0.001; \*\*\*\**p*<0.0001 in a two-way ANOVA test, comparing growth of the clinical isolates to the FGSC-A4 reference strain.

In summary, the aforementioned results suggest strain-specific differences in the response to different physiological stress conditions and infer that the two *A. nidulans* CIs are more sensitive to cell wall-perturbing agents than the reference strain.

### The *A. nidulans* clinical isolates do not display increased resistance to azoles and amphotericin B

Since both CIs showed increased susceptibility to caspofungin, an echinocandin that is being used as a second line treatment for fungal infections (33), and to other cell wall-perturbing agents, we expanded our analyses to include two additional antifungal drugs classes. Specifically, we followed the “Guidelines for the Diagnosis and Management of Aspergillosis”, which, in most of the cases, recommend to treat aspergillosis with azoles and polyene drugs (11), both of which are known to interfere with the biosynthesis or physicochemical properties of fungal membrane sterols (10). Therefore, we determined the minimal inhibitory concentrations (MIC) of the azoles voriconazole, posaconazole and the polyene amphotericin B for all three strains. No differences in the MICs among all strains to these drugs was observed (Table 3).

**Table 3.** Minimum inhibitory concentrations (MIC) of voriconazole, posaconazole and amphotericin B on the *A. nidulans* clinical isolates MO80069 and SP-2605-48 and the FGSC-A4 reference strain.

| Strains | MIC (µg/mL) |  |  |
| --- | --- | --- | --- |
|  | Voriconazole | Posaconazole | Amphotericin B |
| FGSC-A4 | 0.25 | 1.0 | 2.0 |
| MO80026 | 0.25 | 1.0 | 2.0 |
| SP260548 | 0.25 | 1.0 | 2.0 |

### Cleistothecia formation is impaired in the *A. nidulans* SP-2605-48 strain

*A. nidulans* is known for its easily inducible sexual cycle, which serves as a laboratory-based molecular tool for strain construction and studying fungal sexual reproduction (36). To further characterize *A. nidulans* CI biology, we assessed whether *A. nidulans* CIs are able to undergo sexual reproduction, by performing self- and out-crosses for each clinical strain and the reference strain (control) at 30 and 37 °C.

### Strains were first crossed with themselves (self-crosses) at 30 °C and 37

°C, and cleistothecia formation was observed for all strains at both temperatures, except for strain SP-2605-48 at 37 °C (Table 4). Density of cleistothecia (cleistothecia/cm^2^) also varied between strains in a temperature-dependent manner, with the clinical isolates forming fewer cleistothecia per cm^2^ than when compared to the reference strain at 30 °C and 37 °C (Table 4). In addition, no difference in ascospore viability was observed among strains (Table 4).

**Table 4.**
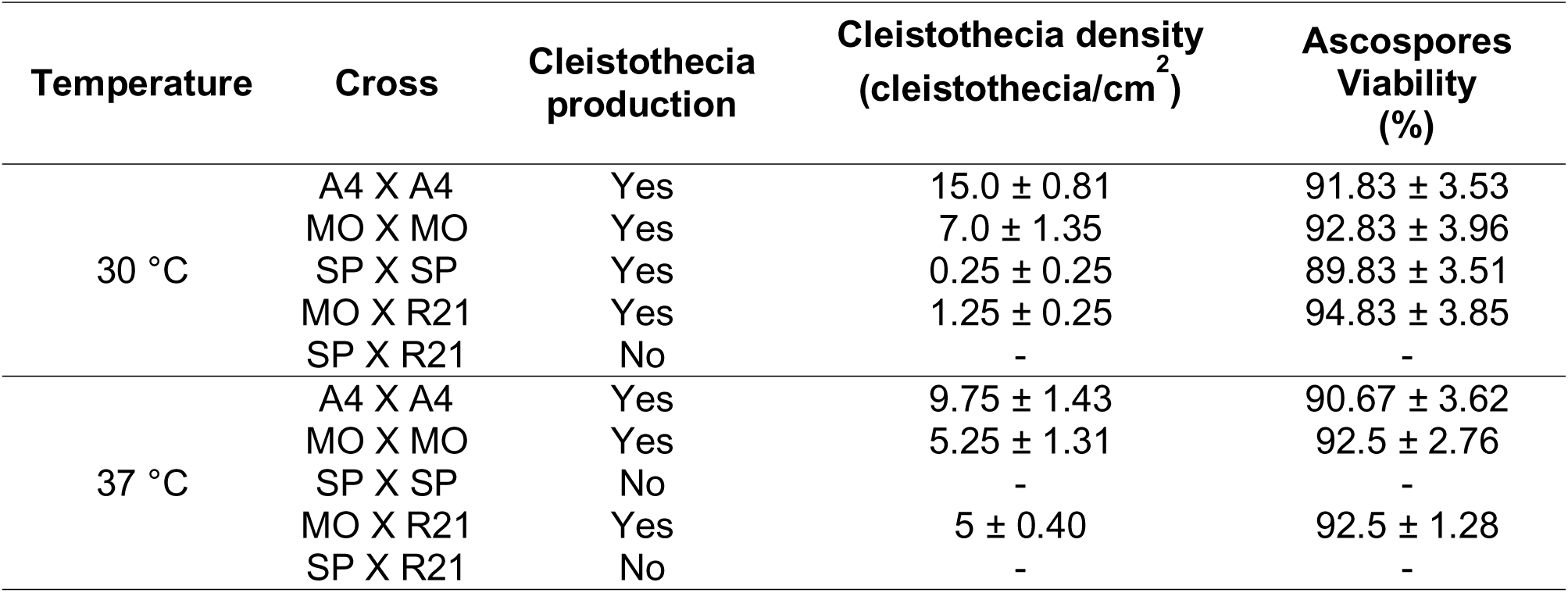
Cleistothecia formation and density and ascospore viability resulting from diverse *A. nidulans* self- and out-crosses (A4 = FGSC-A4 reference strain, MO = MO80069, SP = SP-2605-48).

Out-crosses were performed by crossing the *pyrG* (requirement for uridine and uracil) auxotrophic strains MO80069 and SP-2605-48 with the *paba* (requirement for para-aminobenzoic acid)-deficient strain R21XR135 (Table 6). Strain MO80069 produced cleistothecia at both 30 and 37°C whereas strain SP-2605-48 did not produce any cleistothecia in any of the tested conditions. Density of cleistothecia was very low at 30°C (0.25 cleistothecia/cm^2^) but increased to the same number than observed for the self-crosses at 37°C with high ascospore viability in all cases (Table 4).

### Identification of single nucleotide polymorphisms (SNPs) and copy number variations (CNVs) in the *A. nidulans* clinical isolate genomes

The aforementioned phenotyping and metabolomics results indicate differences between the strains that affect traits such as nutrient source utilization and resistance to different stresses. These results are in agreement with studies in *A. fumigatus* that have described great strain heterogeneity in traits such as growth, fitness and enzyme secretion between different environmental and clinical isolates (24, 37). Indeed, the number of SNPs, obtained during strain pairwise comparison, in the genomes of different *A. fumigatus* strains, range between ~13,500 (24) and ~50,000 (38, 39). Strain heterogeneity has therefore mainly been investigated in environmental and clinical isolates of *A. fumigatus,* whereas similar studies have not been carried out for *A. nidulans* isolates. We therefore decided to determine differences at the genomic level by sequencing the genomes of our two *A. nidulans* CIs and comparing them to the FGSC A4 reference genome.

The genomes of MO80069 and SP-2605-48 aligned at 98.3% and 97.4%, respectively, to the genome of the reference strain FGSC A4 with 99.8% nucleotide identity. On the other hand, 1.5% and 1.9% of the A4 assembled genome did not align to the MO800069 and SP-2605-48 genomes respectively, indicating differences among the genomes of all three strains.

A total of 12,956 and 12,399 SNPs with respect to the A4 reference genome were detected in the genomes of MO80069 and SP-2605-48, respectively (Table 5, Table S2). When comparing the genome of SP-260548 to the genome of MO80069, 12,836 SNPs were detected (Table 5, Table S2). Each SNP mutation was classified as either high, moderate or low, according to their impact on the DNA codon frame and amino acid sequence. High impact type mutations encompass frameshift mutations and stop codon gain/loss, whereas missense mutations, resulting in amino acid changes, are considered as moderate impact-type mutations. Low impact-type mutations contain all synonymous mutations and mutations within gene introns and UTRs (untranslated regions). The genome of MO80069 contained 501 high impact mutations, 6,271 missense (moderate impact) and 6,184 synonymous (low impact) mutations in comparison to the reference genome (Table 5, Table S2). In the genome of SP-2605-48, 465 high impact mutations, 5,896 moderate impact mutations and 6,038 low impact mutations were detected in comparison to the reference genome (Table 5, Table S2). When comparing the genomes of both CIs, 426 high impact mutations, 6,288 missense mutations and 6,122 synonym mutations were detected (Table 5, Table S2). All non-synonymous mutations were distributed throughout the genomes of both CIs and no clear pattern in mutation accumulation could be observed for any of the 8 chromosomes (Fig. 4-5).

**Figure 4.**
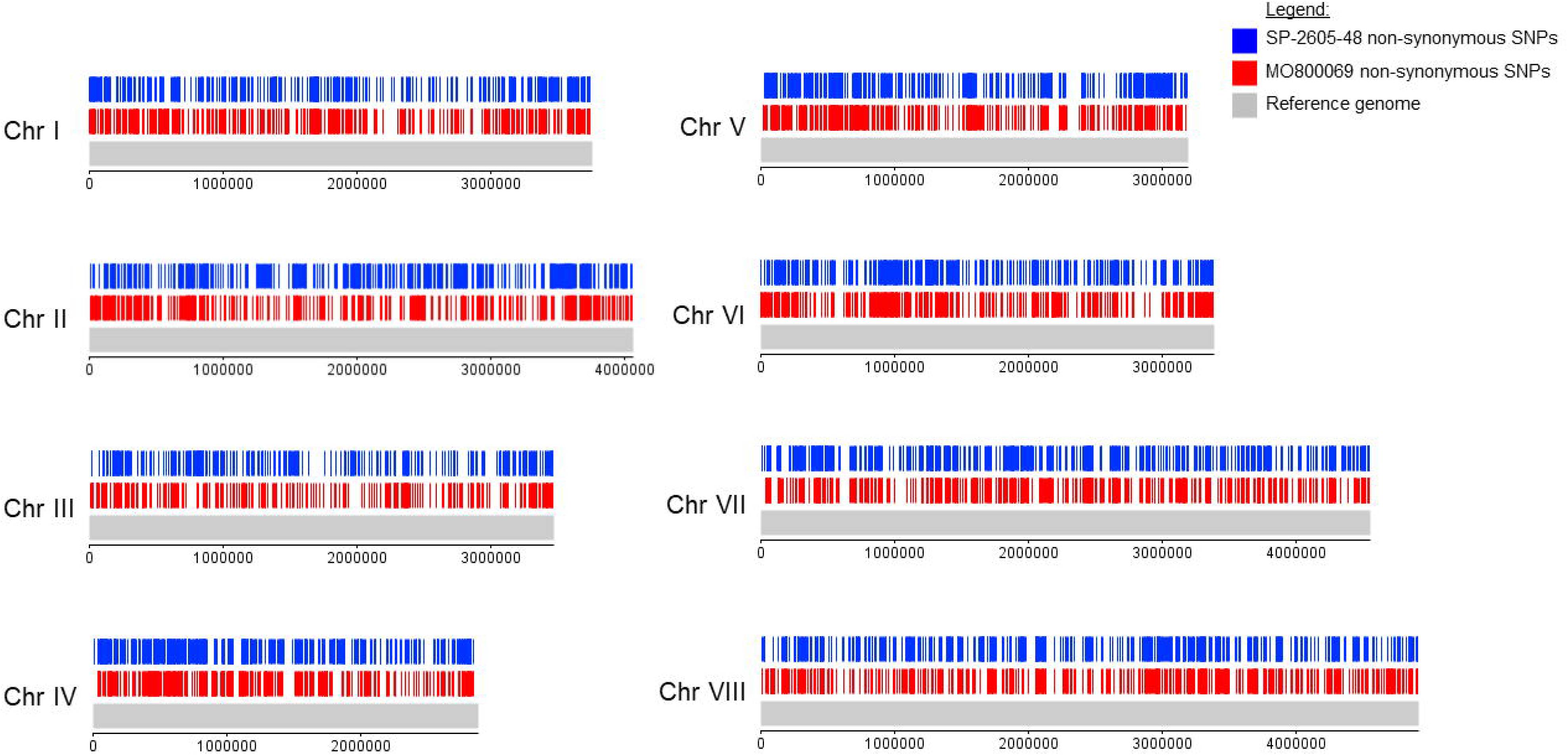
Diagram depicting the location of all detected non-synonymous single nucleotide polymorphisms (SNPs) on the 8 chromosomes (chr I - VIII) of the *A. nidulans* clinical isolates SP-2605-48 and MO80069 in comparison to the FGSC-A4 reference genome.

**Figure 5.**
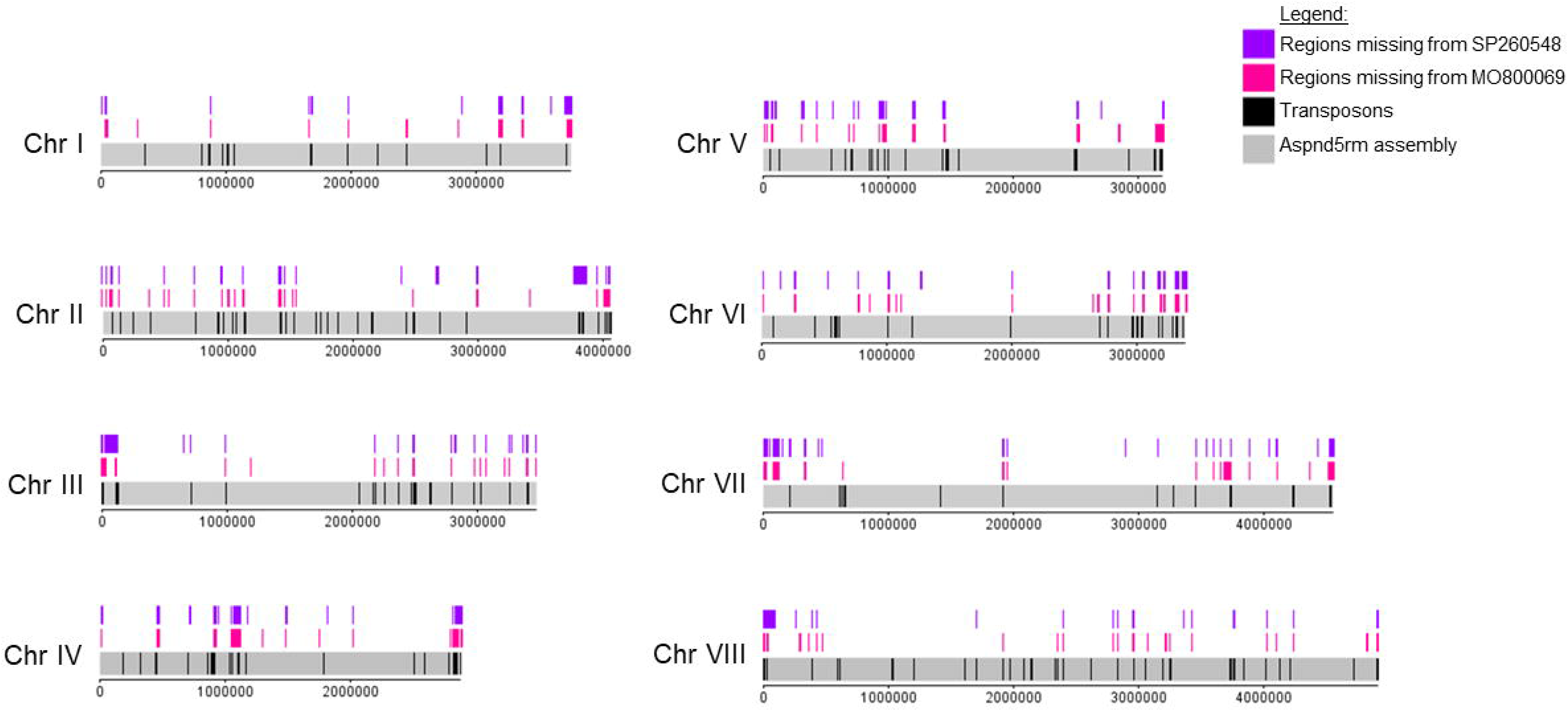
Diagram depicting the location of all detected small deletions on the 8 chromosomes (chr I - VIII) of the *A. nidulans* clinical isolates SP-2605-48 and MO80069 in comparison to the FGSC-A4 reference genome. Also shown are the location of putative transposons in the *A. nidulans* reference genome.

**Table 5.** Type and total amount of single nucleotide polymorphisms (SNPs) and long insertions and deletions (indels) detected in the genomes of the *A. nidulans* clinical isolates MO80069 and SP-2605-48 when compared to the FGSC-A4 reference genome or in both clinical strains.

| Mutation | MO80069 and<br>FGSC A4 | SP-2605-48 and<br>FGSC A4 | SP-2605-48 and<br>MO80069 |
| --- | --- | --- | --- |
| Stop codon gain/loss | 149 | 110 | 170 |
| Frameshift | 352 | 355 | 256 |
| Missense | 6,271 | 5,896 | 6,288 |
| Synonymous | 6,184 | 6,038 | 6,122 |
| **Total SNPs** | **12,956** | **12,399** | **12,836** |
| Insertions | 234 | 308 | 222 |
| Deletion | 114 | 138 | 207 |
| **Total Indels** | **348** | **446** | **375** |

**Table 6.** Strains used in this study (NA = not applicable).

| Strain | Genotype | Source | Reference |
| --- | --- | --- | --- |
| FGSC-A4 | Glasgow Wild type<br>(*veA*+) | Soil | Pontecorvo et al (1953) |
| MO80069 | Wild type, clinical isolate | Bronchoalveolar lavage of a patient with breast carcinoma and pneumonia (Portugal) | This study |
| SP-2605-48 | Wild type, clinical isolate | Patient with cystic fibrosis who underwent lung transplantation (Belgium) | This study |
| R21XR135 | *pabaA1;yA2* | NA | This study |
| MO80069*pyrG*- | *pyrG89* | This study | This study |
| SP-2605-48<br>*pyrG*- | *pyrG89* | This study | This study |
| *ΔmpkA* | *ΔakuB mpkA::ptrA; PTR* | NA | Manfioli et al. (2019) |

In addition, the genomes of both CIs were screened for large-scale (>50 bps) insertions and deletions (indels). In total, 1169 large-scale indels, consisting of anything between 3 bp to 23 kbp in size, were detected on any of the eight chromosomes of the CIs when compared to the reference strain (Supplementary Table 3). Of these, 348 indels were specifically located in the genome of MO80060, 446 indels were found in the genome of SP-2605-48 only, and 375 indels were located in the genomes of both CIs (Table 5, Supplementary Table 3). The majority of these indels were insertions (Table 5). Of the 375 indels found in the genomes of both CIs, 227 (60.5%) indels differed between the two strains, with the remaining 148 indels being identical for both strains (Supplementary Table 3).

### The *A. nidulans* clinical isolates are defect in MpkA accumulation in response to cell wall stress

As this work aimed to characterize metabolic utilization of physiologically-relevant carbon and lipid sources in *A. nidulans* CIs, including acetate and fatty acids, we screened genes encoding proteins important for carbohydrate and lipid utilization, cell wall biosynthesis/remodeling and sexual reproduction for the presence of any of the aforementioned moderate and high impact mutations (Table S4). Moderate impact (missense) mutations were detected in three genes *(hxkA; swoM; pfka),* encoding proteins involved in glycolysis (hexokinase, glucose-6-phosphate isomerase, 6-phosphofructokinase) in both CIs; whereas four and six missense mutations were found in two genes *(idpA* and *mdhA)* encoding the enzymes isocitrate dehydrogenase and malate dehydrogenase of the tricarboxylic acid cycle in the genomes of MO80069 and SP-2605-48, respectively (Table S4). Similarly, several moderate impact mutations were found in genes encoding enzymes required for C2-associated metabolism (acetate, ethanol and fatty acid), including *farA* (transcription factor regulating fatty acid utilization) and *farB* (transcription factor regulating the utilization of short-chain fatty acids) in both CIs, *facA* (acetyl-coA synthase), *acuM* (transcriptional activator required for gluconeogenesis) and *alcM* (required for ethanol utilization) in SP-2605-48 and *echA* (enoyl-coA hydratase) in MO80069 (Table S4). Genes encoding proteins that function in the glyoxylate cycle also contained missense mutations in both CIs (Table S4). Furthermore, a frameshift mutation was detected in both CIs in *acuL,* encoding a mitochondrial carrier involved in the utilization of carbon sources that are metabolized via the Krebs cycle (40) (Table S4). The aforementioned mutations could underlie the observed differences in phenotypic growth in the presence of different carbon and lipid sources.

Due to the absence of cleistothecia formation in strain SP-2605-48, we wondered whether this strain contained any mutations in genes encoding proteins required for *A. nidulans* sexual reproduction. We found 11 and 13 mutations in 7 and 9 genes related to mating in MO80069 and SP-2605-48 genomes, respectively (Table S4). Those mutations include missense and frameshift mutations in genes involved in the perception of light and dark *(ireA, ireB, cryA, veA, velB*), mating processes *(cpcA, rosA, nosA)* and signal transduction *(gprH* and *gprD)* (Table S4). Indeed, *rosA* was absent in both CIs whereas *ireA* was missing from the genome of SP-2605-48. RosA is a transcriptional repressor of sexual development (41) whereas IreA is a transcription factor required for the blue light response, important for developmental processes, including mating.

Lastly, as both Cis were sensitive to cell wall perturbing agents, we screened for mutations in genes encoding enzymes involved in cell wall biosynthesis and degradation. Compared to the FGSC A4 reference genome, we found 159 and 90 mutations in 40 and 34 genes involved in cell wall biosynthesis, integrity and signaling in the genomes of MO80069 and SP-2605-48, respectively (Table S4). The majority of these mutations were moderate impact missense mutations in genes that encode components required for 1,3-β and α-glucan, chitin synthesis and degradation, including various types of glucanases, chitinases and chitin synthases (Table S4). However, 17 (MO80069) and 9 (SP-2605-48) mutations were high impact level mutations which occurred in genes AN0550 (putative glucan 1,3-beta-glucosidase), AN0509 (putative chitinase), AN0517 (putative chitinase), AN0549 (putative chitinase), AN9042 (putative alpha-1,3-glucanase), AN6324 (putative α-amylase), AN4504 (putative endo-mannanase) and AN0383 (putative endo-mannanase) (Table S4). In addition, small frameshift mutations were detected in three genes encoding the mitogen-activated protein kinase (MAPK) kinase kinase BckA (AN4887), the MAPK MpkA (AN5666) and the transcription factor RlmA (AN2984) (Table S4). In *A. fumigatus,* BckA and MpkA are components of the cell wall integrity (CWI) pathway, which ensures the integrity of the cell wall and is activated in response to different cell wall stresses including those exerted by cell wall-targeting anti-fungal drugs (42). RlmA was shown to act downstream of MpkA, regulating cell wall biosynthesis-related genes and this transcription factor is also involved in the direct regulation of MpkA (43). Mutation in *rlmA* was observed only in the genome of strain SP-2648-05.

In order to determine whether the observed frameshift mutations had an impact on CWI signaling, we carried out a western blot of phosphorylated MpkA in the presence of NaCl-induced cell wall stress in all three *A. nidulans* strains. Phosphorylated MpkA levels were normalized by total cellular MpkA. Low levels of phosphorylated MpkA were detected in the absence of NaCl in all three strains, but, whereas MpkA protein levels significantly increased upon cell wall stress in the FGSC A4 reference strain, no phosphorylated MpkA could be detected in both CIs (Fig. 6). These results suggest that the observed frameshift mutations in *mpkA* had an effect on MpkA protein levels in the presence of cell wall stress, potentially being (one of) the cause(s) for the observed increased sensitivity to cell wall-perturbing agents.

**Figure 6.**
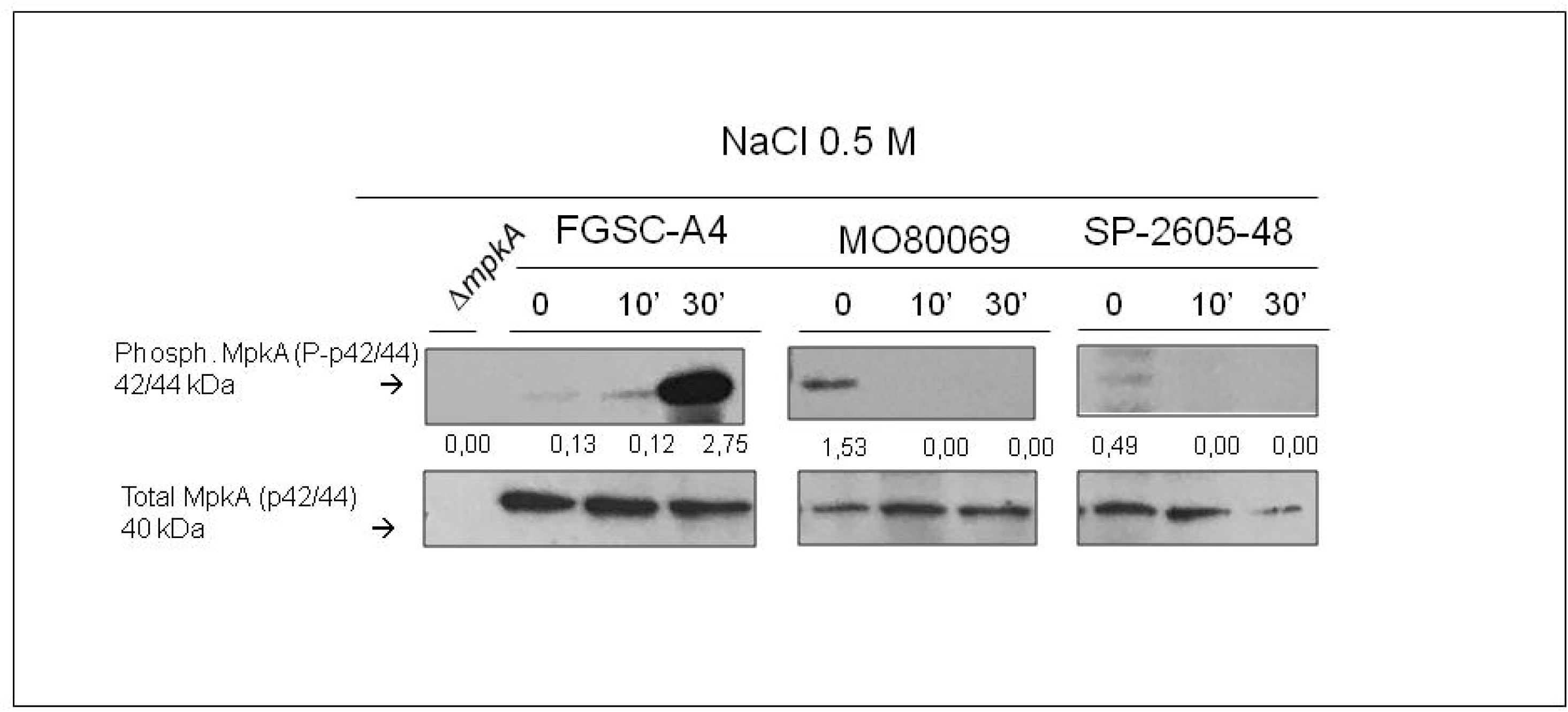
MpkA is not phosphorylated in the *A. nidulans* clinical isolates MO80069 and SP-2605-48 in the presence of NaCl-induced cell wall stress when compared to the FGSC-A4 reference strain. Strains were grown from 1×10^7^spores in complete medium for 16 h (control, 0 min) at 37°C before 0.5 M NaCl was added for 10 and 30 min. Total cellular protein was extracted and western blotting was carried out probing for phosphorylated MpkA. Signals were normalized by the amount of total MpkA present in the protein extracts and cellular extracts from the Β*mpkA* strain were used as a negative control.

### The *A. nidulans* clinical isolates do not display increased resistance to *in* vitro-mediated killing by different types of macrophages and neutrophils

Due to the observed phenotypic and genotypic differences, we wondered whether the CIs were different in virulence from the reference strain. Virulence was first characterized in a variety of *in vitro* conditions. Macrophages play an essential role in clearing *Aspergillus spp* conidia from the lung (8), whereas neutrophils are predicted to primarily be responsible for eliminating fungal hyphae (39). To determine whether any strain-specific differences exist in macrophage-mediated phagocytosis and killing, the respective assays were carried out for all three strains in the presence of murine wild-type and *gp91phox* knockout (CGD) macrophages. Macrophages from CGD patients are impaired in eliminating conidia from the lung environment, thus rendering the host more susceptible to fungal infections (20). Both types of macrophages phagocytised a significantly higher number of conidia from both *A. nidulans* clinical isolates (~75%) when compared to the reference strain (~50%) (Fig. 7A). Indeed, conidia from all three *A. nidulans* strains had increased viability after phagocytosis by *gp91phox* knockout macrophages than when compared to wild-type macrophages, confirming the inability of this type of macrophage to efficiently kill fungal conidia (Fig. 7B). Despite increased phagocytosis of both CIs, no difference in conidial viability was observed for strain MO80069 when compared to the reference strain, whereas wild-type but not CGD macrophages succeeded in killing significantly more SP-2605-48 conidia (Fig. 7B).

**Figure 7.**
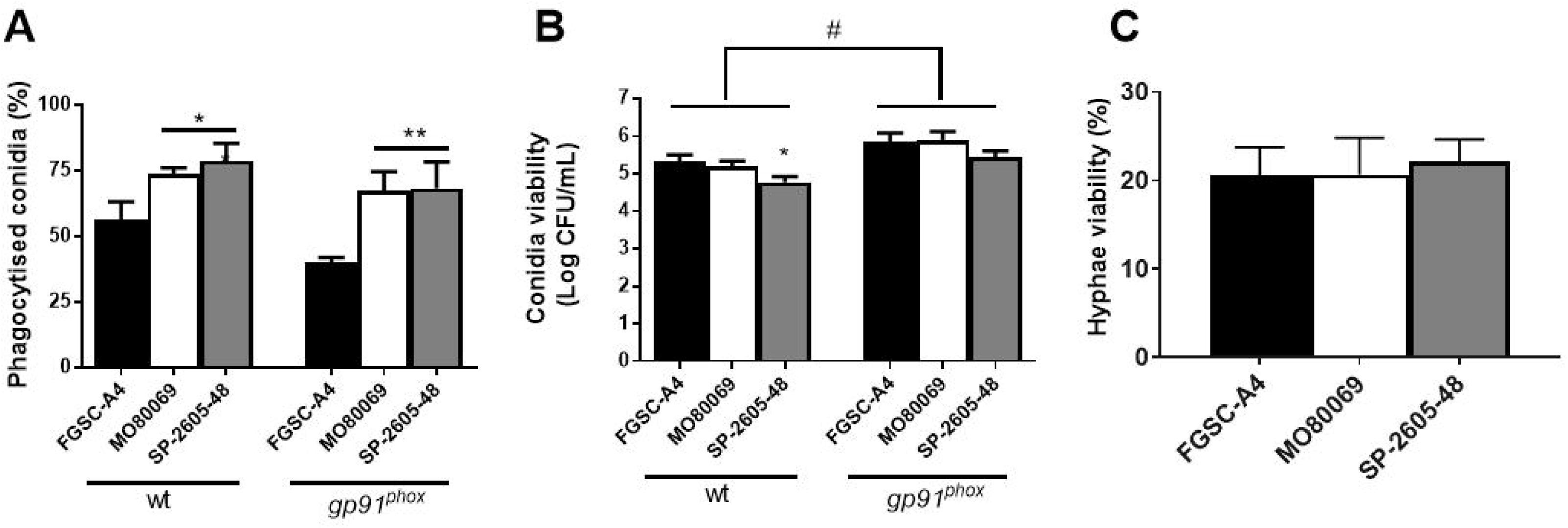
The *A. nidulans* clinical isolates MO80069 and SP-2605-48 do not present increased survival in the presence of macrophages and neutrophils. **(A)** Percentage of phagocytised conidia by murine wild-type and *gp91^phox^* knockout macrophages. Macrophages were incubated for 1.5 h with conidia from the respective strains before phagocytised conidia were counted. **(B)** Colony forming units (CFU) as a measure of conidia viability after passage through wild-type (wt) and *gp91^phox^* knockout macrophages. Macrophages were incubated with the respective conidia for 1.5 h before they were lysed and contents were plated on complete medium. **(C)** Percentage of viable hyphal germlings after incubation for 16 h with neutrophils from healthy human donors. Strain viability was calculated relative to incubation without PMN cells, which was set at 100% for each sample. Standard deviations represent biological triplicates with \**p*<0.05 and \*\**p*<0.01 when comparing the clinical isolates to FGSC-A4; *#p*<0.05 comparing the same strain in the two types of macrophages in a one-way ANOVA test with Tukey post-test.

When challenged with human PMN (polymorphonuclear) cells, fungal survival was reduced approximately 80% for all three *A. nidulans* strains, indicating that the neutrophils were actively killing the hyphal germlings (Fig. 7C). No difference in strain survival was observed for the CIs (Fig. 7C). These results suggest that the *A. nidulans* CIs do not have higher survival rates in the presence of macrophages and neutrophils.

### Virulence of the *A. nidulans* clinical isolates depends on the host immune status

We determined the virulence of both *A. nidulans* CIs in animal models with different immune statuses. As it is well known that *A. fumigatus* strain-specific virulence is highly dependent on the type of host immunosuppression and model (24,37, 43), we sought to determine if this would also be the case for *A. nidulans.* The virulence of *A. nidulans* CIs was assessed in both zebrafish and murine models of pulmonary and invasive aspergillosis. Furthermore, the immune system of each animal was manipulated in order to give rise to either immunocompetent, CGD or neutropenic/neutrophilic models. As with patients, CGD models of both mice (19) and zebrafish (21) are very susceptible to *A. nidulans* infections. In both immunocompetent- and CGD-type zebrafish and mice, no difference in virulence between the *A. nidulans* clinical isolates and the reference strain was observed (Fig. 8A-D). However, the CI MO80069 was significantly more virulent in neutropenic mice and zebrafish with impaired neutrophil function when compared to the reference strain, whereas no difference in virulence was observed for strain SP-2605-48 (Fig. 8E-F). These results suggest that, like in *A. fumigatus, A. nidulans* virulence depends on the strain and the host immune status.

**Figure 8.**
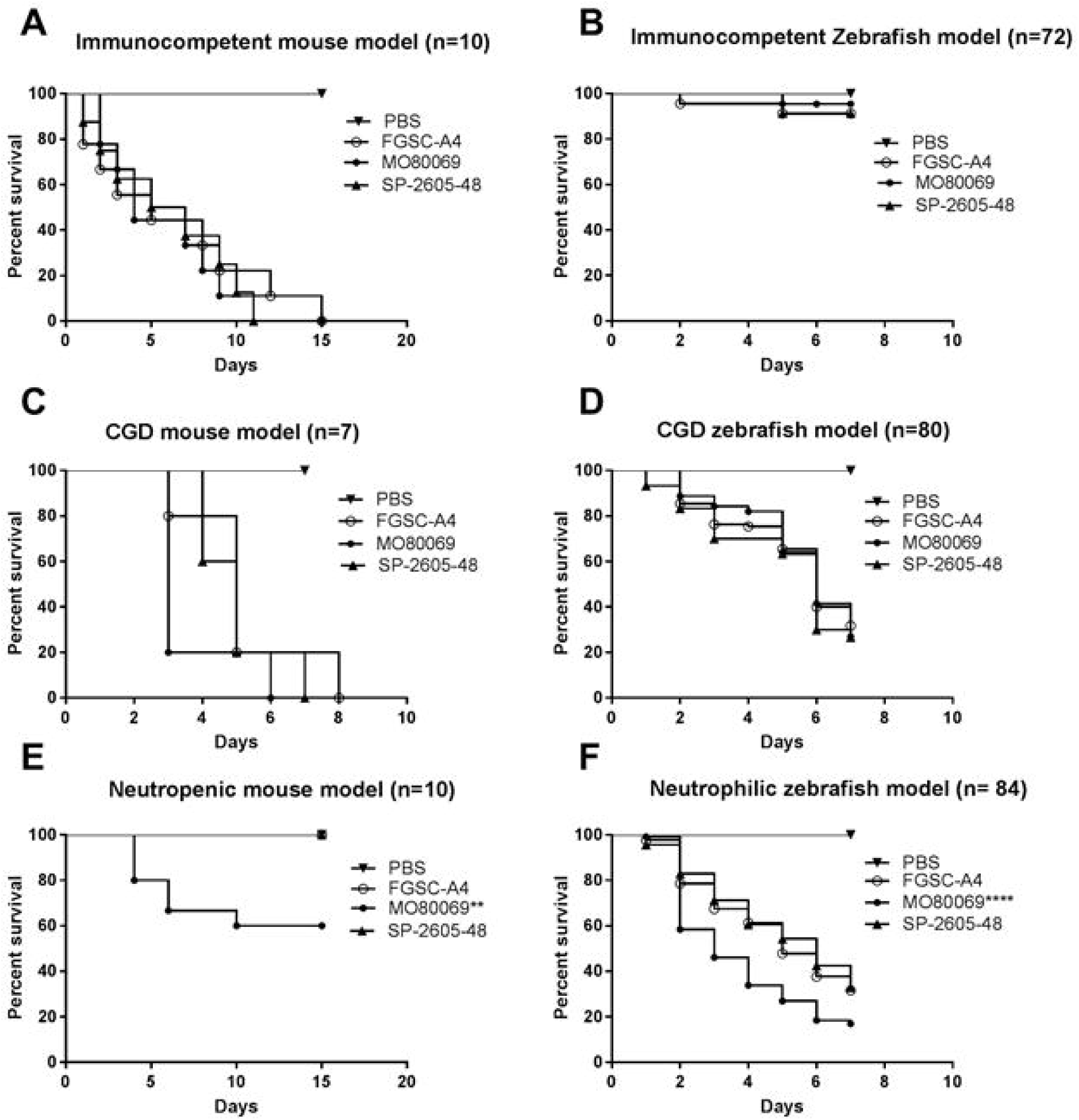
*A. nidulans* strain-specific virulence depends on the host immune status. The virulence of the *A. nidulans* clinical isolates MO80069 and SP-260548 were tested in murine **(A-E)** and zebrafish **(B-F)** models of pulmonary and invasive aspergillosis. Animals were manipulated in order to give rise to either immunocompetent **(A-B),** CGD (chronic granulomatous disease) **(C-D)** or neutropenic (E)/neutrophilic **(F)** models. Shown are survival curves for each immunosuppression and animal model. No difference in virulence was detected for all strains in both immunocompetent and CGD mice. Strain MO80069 was significantly more virulent in neutropenic mice and neutrophilic zebrafish. \*\**p*<0.01; \*\*\*\**p*<0.0001 when comparing survival curves of the clinical isolates to the FGSC-A4 reference strain in a two-way ANOVA test with Tukey post-test.

## Discussion

*Aspergillus nidulans* is a saprophytic fungus that can act as an opportunistic human pathogen in a host immune status- and genetic condition-dependent manner (15,18,44). Infection with *A. nidulans* is prevalent in patients with chronic granulomatous disease (CGD) and isolates have mainly been characterized in the context of this disorder (14, 15). Studies on *A. nidulans* virulence have been carried out in CGD models (animal and cell culture) and virulence characteristics have been compared to the primary human opportunistic fungus *A. fumigatus* (20,21,45,46). *A. fumigatus* infection biology and characterization of strains that were isolated from immunocompromised patients with different conditions have received considerable attention in recent years (24,37,47), whereas similar studies into other pathogenic *Aspergillus spp.* have been neglected, although it is becoming apparent that non-*A. fumigatus* species, including cryptic *Aspergillus* species, also contribute to host infection and invasion (7). This work therefore aimed at providing a detailed phenotypic, metabolic, genomic and virulence characterization of two *A. nidulans* clinical isolates (CIs) that were isolated from non-CGD patients.

The first CI (MO80069) was isolated from a patient with breast carcinoma and pneumonia, whereas the other CI (SP-2605-48) was obtained from a patient with cystic fibrosis who underwent lung transplantation. Genome sequencing confirmed these strains to be *A. nidulans sensu stricto* and growth of these strains was characterized in the presence of physiological-relevant carbon sources. Fungi require carbon sources in large quantities in order to sustain biosynthetic processes and actively scavenge for them in their environment, including mammalian hosts (24). Available carbon sources vary according to the patient’s immune status and disease progression, with, for example, corticosteroid treatment resulting in an increase of fatty and amino acid concentrations and a decrease of glucose levels in mice lungs (22). Growth of the two *A. nidulans* strains in the presence of different carbon sources, differed significantly from the reference strain, with increased biomass accumulation being observed in the presence of alternative (ethanol, lipids, amino acids) carbon sources and reduced growth in the presence of glucose. The observed phenotypic differences were corroborated by metabolic and genomic data which found a number of missense and high impact mutations in genes encoding enzymes required for alternative carbon source and glucose utilization. These included missense mutations in genes encoding glycolysis- and citric acid cycle-related enzymes as well as five missense mutations in the transcription factor-encoding gene *farA,* which regulates the utilization of short- and long-chain fatty acids. Whether these mutations alone and/or in combination with other identified gene mutations are responsible for the observed growth phenotypes remains to be determined. Nevertheless, it is noteworthy that these mutations are found in both CIs, suggesting that these strains are able to grow well in nutrient-poor environments, such as the lung, when compared the reference strain, which was isolated from the soil environment. Furthermore, whether these mutations are a result of adaptation to the host environment also remains subject to future investigations.

In addition, we also assessed the resistance of these strains to a variety of physiological-relevant stress conditions by growing them in the presence of oxidative- and cell wall stress-inducing compounds, high temperature, iron limitation and anti-fungal drugs. Some minor strain-specific differences were observed in these conditions, but the CIs were not significantly more resistant to these conditions in comparison to the reference strain, including azole- and polyene-type anti-fungal drugs. It is possible that the patient-specific lung environment, biofilm formation and/or interactions with other microorganisms may result in protection from or in the absence of these stresses, thus resulting in strains that do not have increased stress tolerance. In contrast to *Candida albicans,* an opportunistic fungal pathogen which was shown to interact with the gram-negative bacterium *Pseudomonas aeruginosa* to promote colonisation of patients with cystic fibrosis in a condition-dependent manner (48), such interactions have not been investigated for *Aspergillus spp. Aspergillus* inter-species interactions in lung microbiomes of patients with and without cystic fibrosis therefore remains an intriguing aspect of fungal pathobiology that warrants further characterization.

In contrast, both *A. nidulans* clinical strains were significantly more sensitive to the cell wall perturbing agents calcofluor white, congo red and caspofungin (33–35) than the reference strain. These results suggest differences in cell wall composition and/or organization between the clinical isolates and the reference strain. When analyzing the respective genome sequences, we found 159 and 90 mutations in 40 and 34 genes encoding enzymes required for cell wall glucan and chitin biosynthesis and degradation in strains MO80069 and SP-2605-48, respectively, when compared to the FGSC-A4 reference strain. Of particular interest was the identification of high impact mutations in genes *bckA, mpkA,* and *rlmA,* which encode components of the CWI signaling pathway. Indeed, Western blotting confirmed the absence of MpkA phosphorylation in the CIs in the presence of cell wall stress. These results suggest that the observed gene mutations cause an altered CWI response, resulting in increased sensitivity to cell wall perturbing agents. The physiological relevance of these findings remains to be determined.

*Aspergillus nidulans* is characterized by an easy inducible sexual cycle as well as by undemanding laboratory-based cultivation and genetic manipulation conditions, and has extensively been used as a model organism to study sexual reproduction and developmental processes (49). Nevertheless, it is unknown whether these traits can also be applied to *A. nidulans* clinical strains and this work therefore assessed the ability of the two CIs to form cleistothecia in self- and out-crosses. Strain MO80069 produced cleistothecia and viable ascospores similar to the reference strain in all tested conditions, whereas strain SP-2605-48 only formed cleistothecia and viable ascospores in self-crosses at 30°C and not 37°C. This suggests that a certain degree of heterogeneity exists with regards to sexual reproduction in *A. nidulans* clinical strains, although a bigger sample size and further studies are required in order to confirm this. Temperature has been shown to influence cleistothecia formation in *Aspergillus spp.* with lower temperatures of 30°C resulting in a higher number of formed cleistothecia (50). Furthermore, we cannot exclude the possibility that strains such as SP-2605-48 may require a different condition for sexual reproduction as it is determined by a series of environmental factors that can either activate or repress sexual development (50). This work identified six missense mutations in four genes *(veA, cpcA, fhbB* and *gprH)* encoding enzymes involved in sexual development and gene *ireA* was absent in the SP-2605-48 genome when compared to strains FGSC-A4 and MO80069. Genes *veA, cpcA, fhbB and ireA* encode proteins that are involved in the perception of environmental signals (50), favouring the hypothesis that SP-2605-48 may require different/specific conditions for cleistothecia production, although it remains to be determined whether the aforementioned mutations and *ireA* are directly linked to the absence of cleistothecia production in strain SP-2605-48 in the conditions tested here.

Lastly, this work examined the *in vivo* virulence of the *A. nidulans* CIs in different animal models with a variety of immune statuses, as *A. fumigatus* strain-specific virulence is highly dependent on the type of host immunosuppression and model (24,37,51). No difference in virulence was observed in immunocompetent and CGD murine and zebrafish models whereas strain MO80069 was significantly more virulent in a zebrafish with impaired neutrophil function and a neutropenic murine model of invasive aspergillosis than when compared to strains FGSC-A4 and SP-2605-48. These results suggest that neutrophil recruitment and function at the site of infection are important for controlling *A. nidulans* infection in both vertebrates. Furthermore, results are in agreement with studies on *A. fumigatus* which show that virulence is as much a strain-dependent as it is a host-dependent trait (24,37,39,51). Furthermore, the tested phenotypes and genome mutations appear to not correlate with strain virulence, although sample size has to be increased in order to confirm this in future studies. *Aspergillus* infection biology of mammalian hosts is a multi-factorial and -faceted process that not only depends on strain-specific virulence traits (30), but also on the genetic composition of the host and status of the immune system (52). Furthermore, the composition and inter-species interactions of the lung microbiome also influences pathogenicity of a given microorganism, with interactions between different species shown to influence host immune responses (49, 53). *A. fumigatus* is the main etiological agent of *Aspergillus-related* diseases and is predominantly present in the lung environment when compared to other infections caused by *Aspergillus spp.* (7). It is therefore possible that other *Aspergillus spp.,* such as *A. nidulans,* remain largely undetected in the lung environment, due to the predominant nature and/or inhibitory function of other fungal species, and where they can grow without the necessity to evolve and adapt to extreme stress conditions. The prevalence and virulence of non-A *fumigatus* species therefore remains a highly interesting and somewhat neglected topic that warrants future detailed studies. In summary, this is the first study that presents extensive phenotypic, metabolic, genomic and virulence characterization of two *A. nidulans* clinical isolates. Just as in *A. fumigatus,* strain heterogeneity exists in *A. nidulans* clinical strains that can define virulence traits. Further studies are required to fully characterize *A. nidulans* strain virulence traits and pathogenicity.

## Materials and Methods

### Ethics statement

The principles that guide our studies are based on the Declaration of Animal Rights ratified by the UNESCO on the 27^th^ January 1978 in its 8^th^ and 14th articles. All protocols used in this study were approved by the local ethics committee for animal experiments from Universidade de São Paulo, Campus Ribeirão Preto (permit number 08.1.1277.53.6). All adult and larval zebrafish procedures were in full compliance with NIH guidelines and approved by the University of Wisconsin-Madison Institutional Animal Care and Use Committee (no. M01570 - 0-02-13).

### Strains, media, and growth conditions

All strains used in this study are listed in Table 6. *A. nidulans* strain FGSC-A4 was used as a reference strain. In addition to culture macroscopic features and fungal microscopic morphology analysis, whole genome sequencing and phylogenetic analysis confirmed that both clinical isolates are *A. nidulans* (Fig S4). For phylogenetic tree construction, we compared *CaM, BenA, RPB2* and *ITS* rDNA sequences, identified using blastN implemented in BLAST+ v2.8.1 (54), to sequences from other species in the *Aspergillus* section *Nidulantes* (55), using a maximum-likelihood tree constructed with MEGA v10.1.1 (56). All strains were maintained in 10% glycerol at −80°C.

Strains were grown either in complete medium or minimal medium as described previously (57). Iron-poor MM was devoid of all iron and supplemented with 200 μM of the iron chelators bathophenanthrolinedisulfonic acid (4,7-diphenyl-1, 10-phenanthrolinedisulfonic acid [BPS]) and 300 μM of 3-(2-pyridyl)-5,6-bis(4-phenylsulfonic acid)-1,2,4-triazine (ferrozine). All growth was carried out at 37 °C for the indicated amounts of time, except where stated. Reagents were obtained from Sigma-Aldrich (St. Louis, MO) except where stated. Radial growth was determined by inoculating plates with 10^5^ spores of each strain and incubation for 5 days before colony diameter was measured. Where required, the oxidative stress-inducing compound menadione or the cell wall perturbing compounds congo red (CR), caspofungin and calcofluor white (CFW) were added in increasing concentrations. All radial growth was expressed as ratios, dividing colony radial diameter (cm) of growth in the stress condition by colony radial diameter in the control (no stress) condition. To determine fungal dry weight, strains were grown from 3 x 10^6^ spores in 30 mL liquid MM supplemented with 1% (w/v) of glucose, acetate, mucin or casamino acid or 1% (v/v) of ethanol, Tween 20 and 80 or olive oil for 48 h (glucose) or 72h (others) at 37 °C, 150 rpm. All liquid and solid growth experiments were carried out in biological triplicates.

Growth in the presence of H_2_O_2_ was carried out as serial dilutions (10^5^ - 10^2^ spores) in liquid CM in 24-well plates for 48h in the presence of different concentrations of H_2_O_2_.

## Metabolite analysis

Metabolome analysis was performed as described previously (58). Briefly, metabolites were extracted from 5 mg of dry-frozen, mycelial powder of four biological replicates. The polar phase was dried and the derivatized sample was analyzed on a Cambi-PAL autosampler (Agilent Technologies GmbH, Waldbronn, Germany) coupled to an Agilent 7890 gas chromatograph coupled to a Leco Pegasus 2 time-of-flight mass spectrometer (LECO, St. Joseph, MI, USA). Chromatograms were exported from the Leco ChromaTOF software v. 3.25 to the R software (www.r-project.org). The Target Search R-package was used for peak detection, retention time alignment, and library matching.

Metabolites were quantified by the peak intensity of a selective mass and normalized by dividing them by the respective sample dry-weight. Principal component analysis was performed using the pcaMethods bioconductor package (59, 60). Pathway enrichment analysis was carried out using MetaboAnalyst (http://www.metaboanalyst.ca/MetaboAnalyst/faces/home.xhtml) (61).

## Determination of minimal inhibitory concentrations (MICs)

MICs of amphotericin B, voriconazole and posaconazole, were determined by growing 10^4^ spores/well in 96-well plates containing 200 μI/well of RPMI and increasing concentrations of the aforementioned compounds, according to the M38 3^rd^ edition protocol elaborated by the Clinical and Laboratory Standards Institute (62).

## Induction of cleistothecia formation

Cleistothecia formation through self-crossing was induced by growing the strains on glucose minimal medium (GMM) plates that were sealed airtight and incubated for 14 days at 30 or 37°C. Plates were scanned for the presence of cleistothecia under a light microscope. To assess ascospore viability, five cleistothecia of each strain were collected, cleaned on 4% w/v agar plates and re-suspended in 100 μl water. Ascospores were counted and 100 ascospores were plated on GMM before colony-forming units (CFU) were determined. Cleistothecia density was determined through counting the number of cleistothecia of a certain area and dividing them by the cm^2^ of the area.

Cleistothecia formation through out-crossing was carried out as described previously (57). To induce *pyrG^-^* auxotrophy in strains MO80069 and SP-2605-48 (Table 1), they were grown on GMM plates supplemented with 1.2 g/L uridine and uracil (UU) and 0.75 mg/mL 5-fluoroorotic acid (FOA) in the form of a cross until single colonies appeared. Auxotrophy was confirmed by growing strains on GMM with and without UU before strains were crossed with strain R21XR135 (Table 1).

### DNA extraction, genome sequence, detection of single nucleotide polymorphisms (SNPs), insertions and deletions (lndels)

DNA was extracted as described previously (57). Genomes were sequenced using 150-bp Illumina paired-end sequence reads at the Genomic Services Lab of Hudson Alpha (Huntsville, Alabama, USA). Genomic libraries were constructed with the Illumina TruSeq library kit and sequenced on an Illumina HiSeq 2500 sequencer. Samples were sequenced at greater than 180X coverage or depth. Short-read sequences for these strains are available in the NCBI Sequence Read Archive (SRA) under accession number.

The Illumina reads were processed with the BBDuk and Tadpole programs of BBMap release 37.34 [https://sourceforge.net/projects/bbmap/files/BBMap_37.34.tar.gz/download] to remove sequencing adapters and phiX, and to correct read errors. The *Aspergillus nidulans* FGSC_A4 genome sequence and gene predictions, version s10-m04-r15, were obtained from the Aspergillus Genome Database [http://aspgd.org/]. The processed DNA reads were mapped to the FGSC_A4 genome with minimap2 version 2.17 [https://github.com/lh3/minimap2] and variants from the FGSC_A4 sequence were called with Pilon version 1.23 [https://github.com/broadinstitute/pilon]. Short indels and nucleotide polymorphisms were recovered from the Pilon VCF files by filtering with vcffilter [https://github.com/vcflib/vcflib] to retain only calls with read coverage deeper than 7, exactly one alternative allele, and alternative allele fraction at least 0.8. Longer indels and sequence polymorphisms were recovered by searching the VCF files for the SVTYPE keyword. Sequence variations inside predicted genes and their effects on predicted protein sequence were identified with a custom Python script. The mitochondrial genome was obtained from the discarded contigs of MaSuRCA. Due to its circular nature, the mitochondrial genome appeared repeated multiple times in a single contig. Lastal (http://last.cbrc.jp/doc/last.html) was used to extract one single copy of the mitochondrial genome using the reference mitochondrion.

### Detection of large genome deletions and insertions

Genome assemblies of the two clinical isolates were aligned to the FGSC A4 reference genome with nucmer (Kurtz et al., 2004). The alignments were filtered to keep only one-to-one matches. Strain-specific loci were detected by searching the alignment coordinates table for regions of the A4 genome with no match in the clinical isolate genome. Large insertions were detected by searching the alignment coordinates table for regions of the clinical isolate genomes with no match in the A4 genome.

### Identification of transposon-like regions in the FGSC-A4 reference genome

Transposon-like regions were identified by running Pfam (64) on the six translation frames of the complete genome sequence. Regions containing any of the fourteen domains typically known to be associated with transposable elements (Table S1) were collected. Inverted repeats longer than 50 bp and separated by less than 5000 bp were extracted and marked as potential Miniature Inverted-repeat Transposable elements (MITE). The Pfam and MITE locations were combined to form the transposon track.

### Figure generation

DNAPlotter (65) was used to display the loci of all non-synonymous SNPs and large deletions identified in the two clinical strains when compared to the reference genome of FGSC A4. In addition, the locations of transposon-like regions in the A4 genome are also highlighted using DNAPlotter.

### Western blotting

Strains were grown from 1×10^7^ spores at 37 °C, 200 rpm, in 50 ml CM for 16 h before being exposed to 0.5 M NaCl for 0, 10 and 30 min. Total cellular proteins were extracted according to Fortwendel and colleagues (2010)(66) and quantified according to Hartree and colleagues (1972) (67).

For each sample, 60 μg of total intracellular protein were run on a 12% (w/v) SDS-PAGE gel before they were transferred to a polyvinylidene difluoride (PVDF) membrane (GE Healthcare). Phosphorylated MpkA or total MpkA was probed for by incubating the membrane with a 1:5000 dilution of the anti-phospho-p44/42 MAPK (9101; Cell Signaling Technologies) antibody or with a 1:5000 dilution of the p44-42 MAPK (Cell Signaling Technology) antibody overnight at 4 °C with shaking. Subsequently, membranes were washed 3 x with TBS-T (2.423 g/l Tris, 8 g/L NaCl, 1 ml /l Tween 20), incubated with a 1:5000 dilution of an anti-rabbit IgG horseradish peroxidase (HRP) antibody# 7074 (Cell Signaling Technologies) for 1 h at room temperature. MpkA was detected by chemoluminescence using the Western ECL Prime (GE Healthcare) blot detection kit according to the manufacturer’s instructions. Films were submitted to densitometric analysis using the ImageJ software (http://rsbweb.nih.gov/ij/index.html). The amount of phosphorylated MpkA was normalized by total MpkA. The *A. fumigatus Δmpka* strain was used as a negative control (Table 1) (68).

### Isolation and differentiation of bone marrow-derived murine macrophages

Bone marrow-derived macrophages (BMDMs) were isolated as described previously (69). Briefly, BMDMs were recovered from femurs of C57BL6 wild-type and *gp91^phox^ knockout* mice and were incubated in BMDM medium [RPMI medium (Gibco) supplemented with 30% (v/v) L929 growth conditioning media, 20% inactivated fetal bovine serum (FBS) (Gibco), 2 mM glutamine and 100 units/mL of penicillin-streptomycin (Life Technologies)]. After 4 days, fresh media was added for an additional 3 days before BMDMs were collected.

### *In vitro* phagocytosis and killing assays

Phagocytosis and killing assays of *A. nidulans* conidia by wild-type and *gp91phox knockout* macrophages were carried out according to Bom et al. (2015) (70) with modifications. 24-well plates containing a 15-mm-diameter coverslip in each well (phagocytosis assay) or without any coverslip (killing assay) and 2 x 10^5^ macrophages per well were incubated in 1 ml of RPMI-FBS [(RPMI medium (Gibco) supplemented with 10% inactivated fetal bovine serum (FBS) (Gibco), 2 mM glutamine and 100 units/mL of penicillin-streptomycin (Life Technologies)] at 37 °C, 5% CO2 for 24 h. Wells were washed with 1 ml of PBS before the same volume of RPMI-FBS medium supplemented with 1 x 10^6^ conidia (1:5 macrophage/conidium ratio) was added in the same conditions.

To determine phagocytosis, macrophages were incubated with conidia for 1.5 h before the supernatant was removed and 500 μl of PBS containing 3.7% formaldehyde was added for 15 min at room temperature (RT). Sample coverslips were washed with 1 ml of ultrapure water and incubated for 20 min with 500 μl of 0.1 mg/ml CFW (calcufluor white) to stain for the cell wall of non-phagocytised conidia. Samples were washed and coverslips were viewed under a Zeiss Observer Z1 fluorescence microscope. In total, 100 conidia were counted per sample and the phagocytosis index was calculated. Experiments were performed in biological triplicates.

To determine macrophage-induced killing of conidia, macrophages were incubated with conidia for 1.5 h before cell culture supernatants were collected and cytokine concentrations were determined. Macrophages were then washed twice with PBS to remove all non-adherent cells and subsequently lysed with 250 μL of 3% v/v Triton X-100 for 10 min at RT. Serial dilutions of lysed samples were performed in sterile PBS and plated onto CM and incubated at 37 °C for 2 days, before colony forming units (CFU) were determined.

### Polymorphonuclear (PMN) cell isolation and spore germination assay

Human PMN cells from fresh venous blood of healthy adult volunteers were isolated according to Drewniak et al. (2013) (71), with modifications. Cells were harvested by centrifugation in isotonic Percell, lysed, and re-suspended in 4-(2-hydroxyethyl)-1-piperazineethanesulfonic acid-buffered saline solution. *A. nidulans* asexual spores were incubated with PMN cells (1 x 10^5^ cells/mL; effector: 1:500) in a 96-well plate overnight at 37 °C in RPMI 1640 medium containing glutamine and 10% fetal calf serum (Life). PMN cells were lysed in a water and a sodium hydroxide (pH 11.0) solution (Sigma-Aldrich) and spore germination was determined using an MTT (thiazolyl blue; Sigma-Aldrich) assay, according to Dos Reis et al., (2011). Strain viability was calculated relative to incubation without PMN cells, which was set at 100% for each sample. The viability of *A. nidulans* germinated spores in the presence of PMN cells, was determined as described previously (32).

### *In vivo* immunocompetent, CGD (chronic granulomatous disease) and neutrophilic zebrafish infections

We evaluated strain virulence in an established zebrafish-aspergillosis model (72 Wild-type larvae were used as an immunocompetent model. Larvae with a dominant negative Rac2D57N mutation in neutrophils *(mpx:rac2D57N)* (Rosowski et al., 2017) were used as a model of leukocyte adhesion deficiency, where neutrophils do not reach the site of infection, and p22^phox^_deficient larvae *(p22^phox (sa11798^)* were used as a CGD model (21).

Spore preparation and conidium micro-injection into the hindbrain of 2-days post fertilization (dpf) larvae were performed as previously described (72). Briefly, after manual dechorionation of embryos, 3 nL of inoculum or PBS-control were injected into the hindbrain ventricle via the optic vesicle (~50 conidia) in anesthetized larvae at approximately 36 h post fertilization.

### *In vivo* immunocompetent, CGD (chronic granulomatous disease) and neutropenic murine infections

Virulence of the *A. nidulans* strains was determined in immunocompetent, CGD and neutropenic mice. *A. nidulans* conidial suspensions were prepared and viability experiments carried out as described previously (70). Eight to twelve weeks old wild-type (n=10) and *gp91^phox^ knockout* (n=7) C57BL/6 male mice were used as immunocompetent and CGD models, respectively. Neutropenia was induced in 7-8 weeks old BALB/c female mice (n=10, weighing between 20 and 22 g) with cyclophosphamide at a concentration of 150 mg per kg, administered intraperitoneally (i.p) on days -4 and -1 prior to infection (day 0) and 2 days post-infection. Hydrocortisone acetate (200 mg/kg) was injected subcutaneously on day -3 prior to infection.

Mice were anesthetized and submitted to intratracheal (i.t.) infection as previously described (73) with some minor modifications. Briefly, after i.p. injection of ketamine and xylazine, animals were infected with 5.0 x 10^7^ (immunocompetent) or 1 x 10^6^ (CGD) conidia contained in 75 μL of PBS (74) by surgical i.t. (intratracheal) inoculation, which allowed dispensing of the fungal conidia directly into the lungs. Neutropenic mice were infected by intranasal instillation of 1.0 x 10^4^ conidia as described previously (70). PBS (phosphate buffered saline) was administered as a negative control for each murine model.

Mice were weighed every 24 h from the day of infection and visually inspected twice daily. The endpoint for survival experimentation was identified when a 20% reduction in body weight was recorded, at which time the mice were sacrificed.

### Statistical analyses

All statistical analyses were performed using GraphPad Prism version 7.00 (GraphPad Software, San Diego, CA, USA), with P < 0.05 considered significantly different. A two-way analysis of variance (ANOVA) was carried out on all stress response tests; whereas a one-way ANOVA with Tukey post-test was applied for growth in the presence of different carbon sources, phagocytosis index and PMN cell killing assay. Survival curves were plotted by Kaplan-Meier analysis and results were analyzed using the log rank test. All experiments were repeated at least twice.

## Acknowledgements

We would like to thank the Fundação de Amparo à Pesquisa do Estado de São Paulo (FAPESP, São Paulo research foundation), grant numbers 2016/07870-9, 2017/19821-5, LNAR 2017/14159-2, FVL 2018/14762-3 and 2019/00631-7 and the Conselho Nacional de Desenvolvimento Científico e Tecnológico (CNPq) for financial support. JLS and AR were supported by the Howard Hughes Medical Institute through the James H. Gilliam Fellowships for Advanced Study program. FR was supported by the Northern Portugal Regional Operational Programme (NORTE 2020), under the Portugal 2020 Partnership Agreement, through the European Regional Development Fund (FEDER) (NORTE-01-0145-FEDER-000013).

We also thank Dra. Danielle da Glória de Souza (UFMG-Brazil) for helping with *gp91^phox^ knockout* C57BL/6 mice.

The funders had no role in study design, data collection and interpretation, or the decision to submit the work for publication.

