## Supplementary Figures for "Functional characterization of clinical isolates of the opportunistic fungal pathogen *Aspergillus nidulans*"

**Figure S1. The clinical isolates MO80069 and SP-2605-48 are metabolically different from the FGSC-A4 reference strain.** Principal component analysis (PCA) of the levels of quantified metabolites identified in biological replicates of all three strains when grown for 16 h in minimal medium supplemented with glucose (**A**), ethanol (**B**), acetate (**C**) and mucin (**D**).

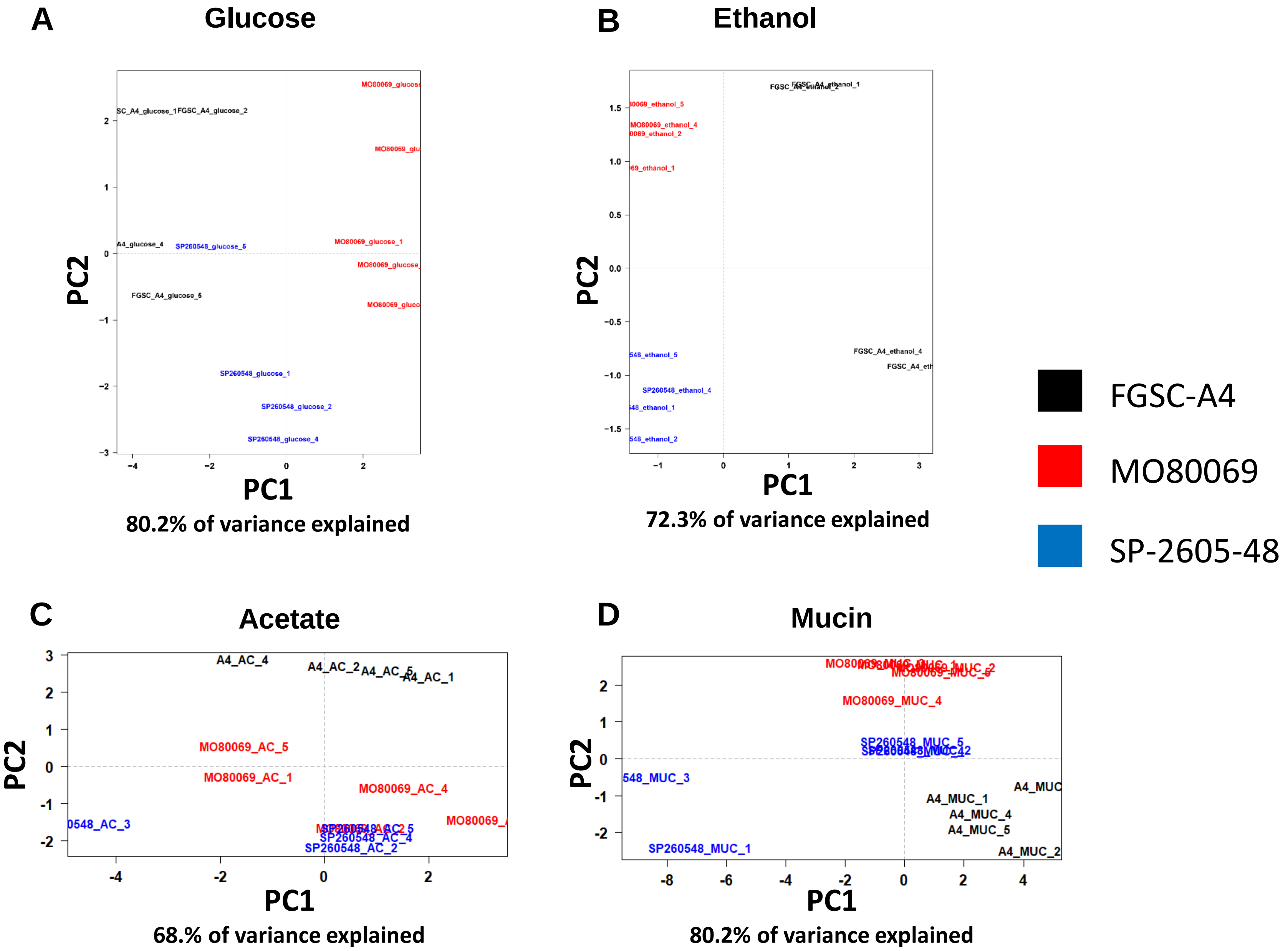

**Figure S2. The clinical isolates MO80069 and SP-2605-48 are metabolically different from the FGSC-A4 reference strain.** Hierarchical component analysis (HCA) of the levels of quantified metabolites identified in biological replicates of all three strains when grown for 16 h in minimal medium supplemented with glucose (**A**), ethanol (**B**), acetate (**C**) and mucin (**D**).

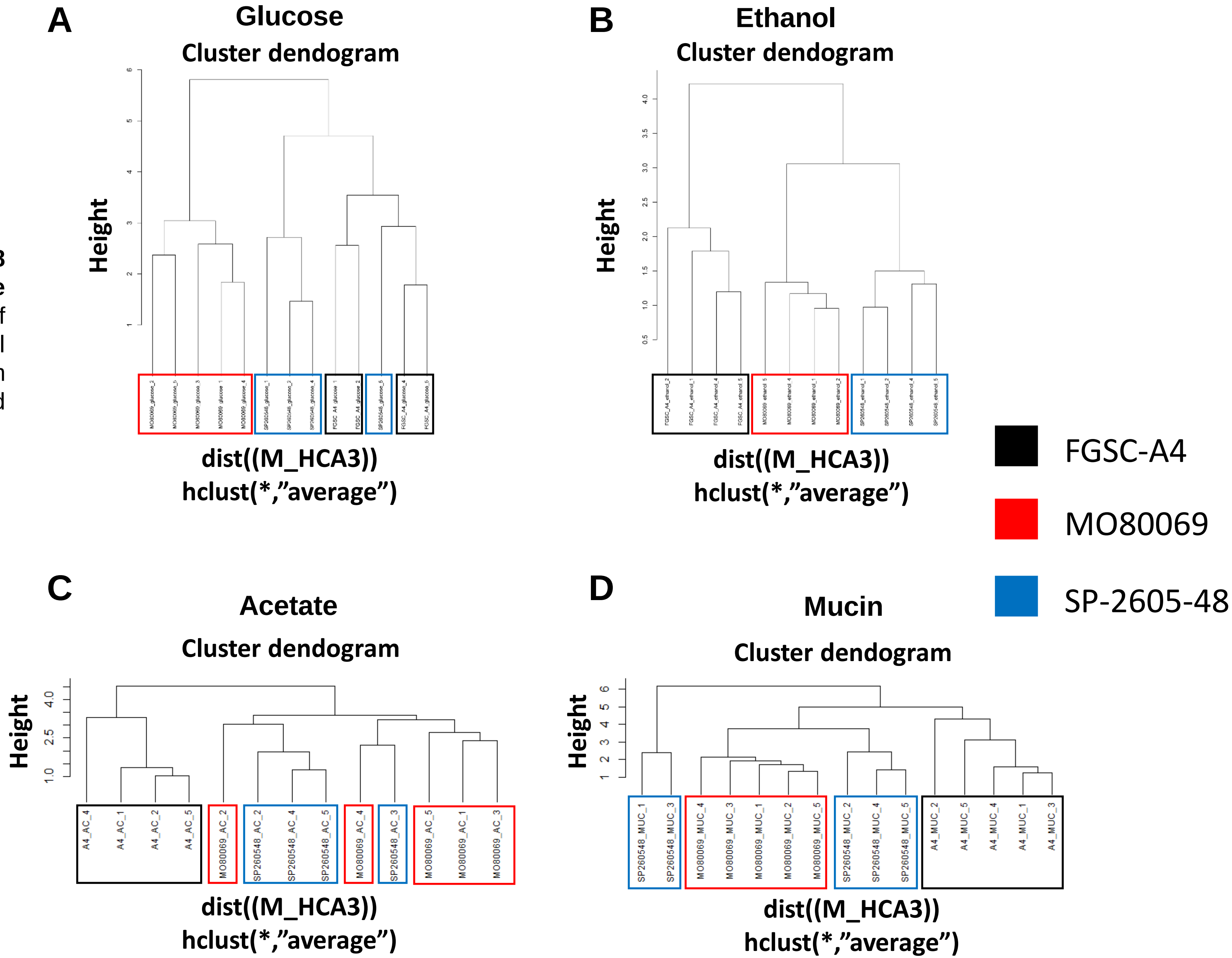

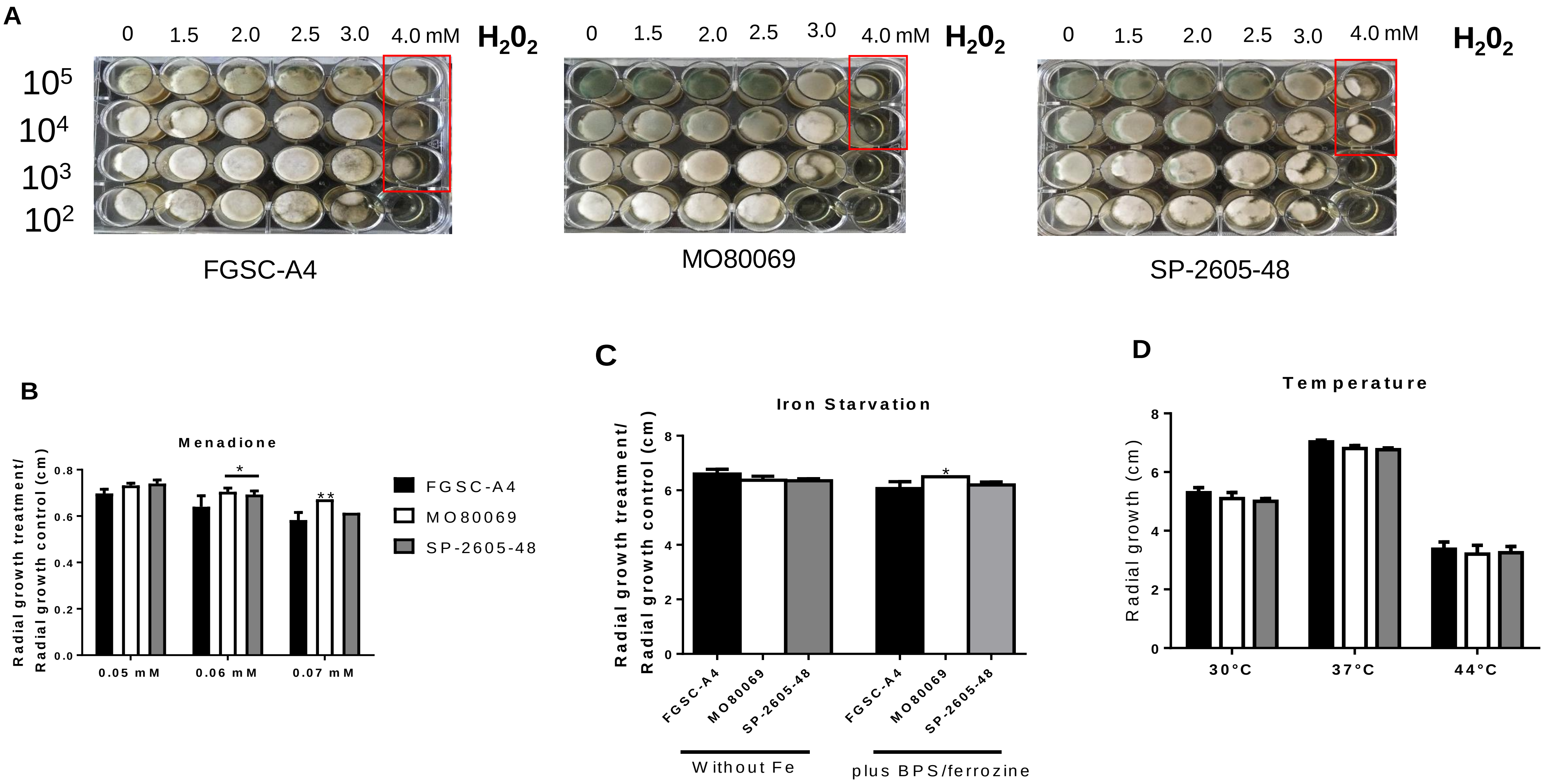


**Figure S3**. **The *A. nidulans* clinical isolates are sensitive to hydrogen peroxide-induced oxidative stress but not to other physiological-relevant stresses**. Strains were grown in tenfold serial dilutions in complete medium supplemented with increasing concentrations of $H_2O_2$ for 48 h at 37°C (**A**), or they were grown from $10^5$ spores on glucose minimal medium (GMM) supplemented with increasing concentrations of menadione (**B**), in the presence of iron-depleted GMM supplemented with the iron chelators BPS and ferrozine (**C**) and on GMM in the presence of different temperatures (**D**) for 5 days at 37°C. Standard deviations represent biological triplicates with *$p<0.05$;**$p<0.01$; ***$p<0.001$; ****$p<0.0001$ in a 2-way ANOVA test when comparing growth of the clinical isolates to the FGSC-A4 reference strain.

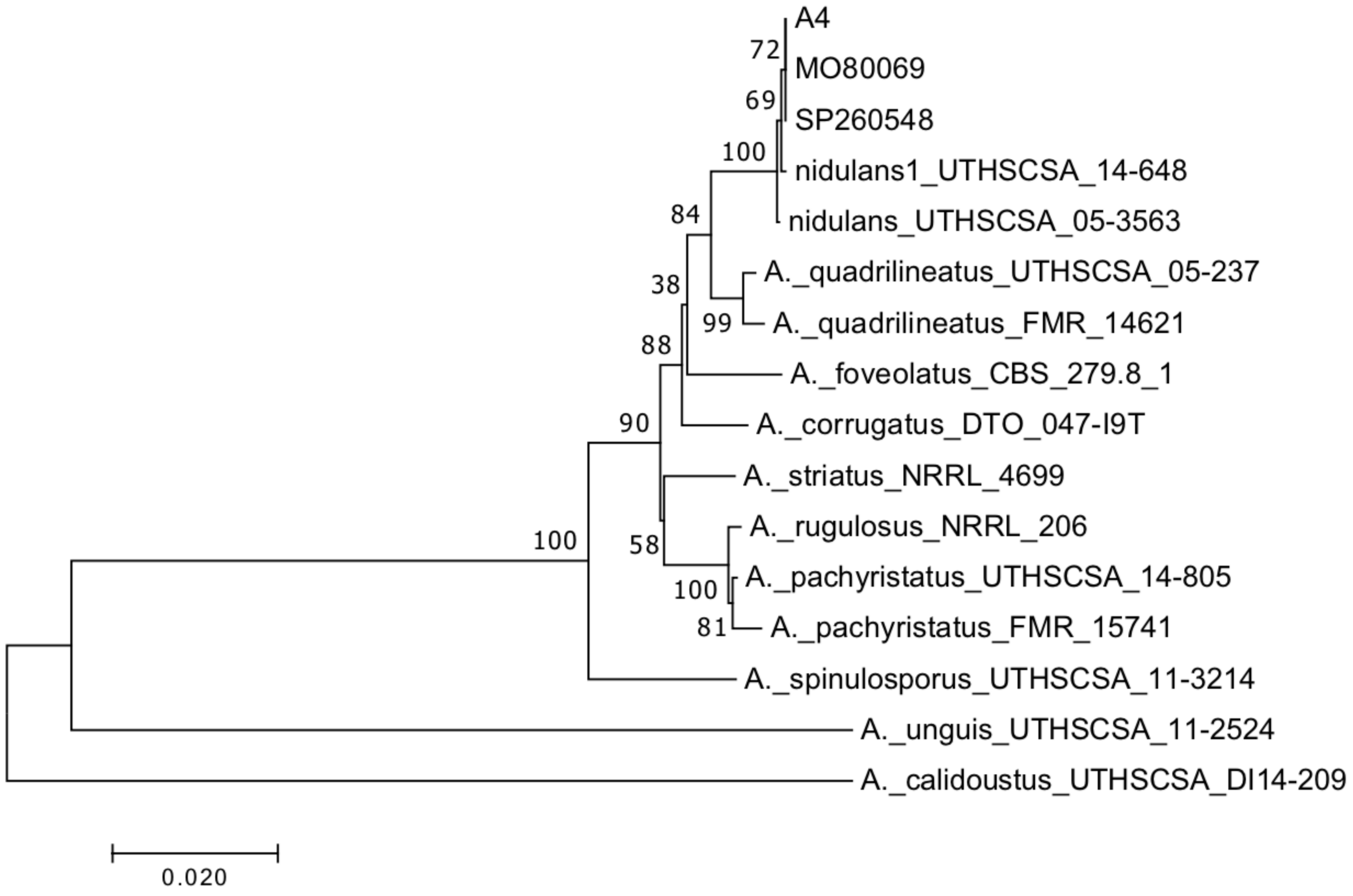


**Figure S4**. **The clinical isolates MO80069 and SP-2605-48 are true *A. nidulans* strains**. The phylogenetic tree was constructed based on the *CaM*, *BenA*, *RPB2* and *ITS* rDNA sequencing data, which was compared to sequences from other species in the *Aspergillus* section of *Nidulantes.*
